# Viscoelastic properties of suspended cells measured with shear flow deformation cytometry

**DOI:** 10.1101/2022.01.11.475843

**Authors:** Richard Gerum, Elham Mirzahossein, Mar Eroles, Jennifer Elsterer, Astrid Mainka, Andreas Bauer, Selina Sonntag, Alexander Winterl, Johannes Bartl, Lena Fischer, Shada Abuhattum, Ruchi Goswami, Salvatore Girardo, Jochen Guck, Stefan Schrüfer, Nadine Ströhlein, Mojtaba Nosratlo, Harald Herrmann, Dorothea Schultheis, Felix Rico, Sebastian Müller, Stephan Gekle, Ben Fabry

## Abstract

Numerous cell functions are accompanied by phenotypic changes in viscoelastic properties, and measuring them can help elucidate higher-level cellular functions in health and disease. We present a high-throughput, simple and low-cost microfluidic method for quantitatively measuring the elastic (storage) and viscous (loss) modulus of individual cells. Cells are suspended in a high-viscosity fluid and are pumped with high pressure through a 5.8 cm long and 200 µm wide microfluidic channel. The fluid shear stress induces large, near ellipsoidal cell deformations. In addition, the flow profile in the channel causes the cells to rotate in a tank-treading manner. From the cell deformation and tank treading frequency, we extract the frequency-dependent viscoelastic cell properties based on a theoretical framework developed by R. Roscoe^1^ that describes the deformation of a viscoelastic sphere in a viscous fluid under steady laminar flow. We confirm the accuracy of the method using atomic force microscopy-calibrated polyacrylamide beads and cells. Our measurements demonstrate that suspended cells exhibit power-law, soft glassy rheological behavior that is cell cycle-dependent and mediated by the physical interplay between the actin filament and intermediate filament networks.

## Introduction

Eukariotic cells can carry out complex mechanical tasks such as cell division, adhesion, migration, invasion, and force generation. These mechanical activities in turn are essential for higher-order cell functions including differentiation, morphogenesis, wound healing, or inflammatory responses. Since cell mechanical activities are accompanied by phenotypic changes in the cell’s viscoelastic properties, measuring them can help elucidate higher-order cell functions in health and disease^2^.

In this report, we describe a quantitative, low-cost, high-throughput, and simple method to measure the viscoelastic properties of cells. The cells are suspended in a high-viscosity (0.5–10 Pa s) fluid (e.g. a 2% alginate solution) and are pumped at pressures of typically between 50–300 kPa through a several centimeter long microfluidic channel with a square cross section (200×200 µm in our set-up). The fluid shear stress induces large cell deformations that are imaged using a complementary metal-oxide-semiconductor (CMOS) camera at frame rates of up to 500 frames/s to achieve a measurement throughput of up to 100 cells/s. Images are stored and analyzed off-line at a speed of around 50 frames/s on a standard desktop PC equipped with a graphics card.

The method takes advantage of two physical principles: First, the shear stress profile inside a long microfluidic channel depends only on the pressure gradient along the channel, which can be precisely controlled, and the channel geometry, which is fixed. Importantly, the shear stress profile does not depend on the viscosity of the cell suspension medium and smoothly increases from zero at the channel center to a maximum value at the channel walls. Accordingly, cells appear circular near the channel center and become increasingly elongated near the channel walls. As the width of the channel is significantly larger than the cell diameter, fluid shear stresses remain approximately constant across the cell surface. From the stress-strain relationship, we estimate the storage modulus of the cell, which characterizes its elastic behavior.

Second, depending on the flow speed profile inside the channel, the cells rotate in a tank-treading manner, similar to a ball that is compressed between two counter-moving parallel plates. The rotational speed of this tank-treading motion increases with increasing shear rate near the channel walls. Tank-treading in combination with the cell’s viscous properties leads to energy dissipation, which limits the increase of cell strain at higher stresses near the channel walls. From this behavior, we extract the loss modulus of the cell, which characterizes its viscous behavior.

For the quantification of cell mechanical properties, we use a theoretical framework developed by R. Roscoe^1^ that describes the deformation of a viscoelastic sphere in a viscous fluid under steady shear flow. This theory allows us to compute the stiffness (shear modulus) and viscosity of a cell from 5 measurable parameters. First, the fluid shear stress acting on the cell must be known, which we compute based on the extension of Poiseuille’s equation to channels with square cross section^3^. Second, we measure the cell deformation (cell strain) from bright-field microscopy images. Third, we measure the alignment angle of the deformed cell with respect to the flow direction. This alignment angle depends on the ratio between cell viscosity and the viscosity of the suspension fluid. Fourth, we compute the local viscosity of the suspension fluid based on measurements of the radial flow speed profile in the channel, which we obtain from multiple images of the same cell during its passage through the channel. Fifth, since cell stiffness and cell viscosity are frequency-dependent, we measure the tank-treading frequency of each cell. To compare the stiffness and viscosity of cells that have experienced different tank-treading frequencies, we scale the stiffness and fluidity of each cell to a reference frequency of 1 rad/s.

Using cell lines and calibrated polyacrylamide beads, we verify that our method provides accurate quantitative measurements of viscoelastic properties. Measurement results are not or only marginally influenced by experimental details such as the viscosity of the suspension fluid or the time point after suspending the cells. We demonstrate that the cell’s viscoelastic properties measured with our method conform to soft glassy power-law rheology that has been reported for a wide range of cells measured with different methods. We also show that our method can be used for dose-response measurements of drugs that induce actin cytoskeleton disassembly, and that these responses are modulated by the cell cycle and the intermediate filament network of the cells.

## Results

### Measurement setup

We image the cells in bright-field mode while they are moving through the microchannel (Fig. 1a-c). Using a neural network, we detect cells that are in focus at the mid-plane of the microchannel (Fig. 1b), and segment their shapes (Fig. 1d). We then quantify the cell position and cell shape by fitting an ellipse to the segmented cell image, from which we obtain the centroid coordinate (*x*_0_, *y*_0_), the length of the semi-major axis *a* and the semi-minor axis *b*, and the angular orientation *β* of the major axis with respect to the *x*-(flow) direction (Fig. 1e). From *a* and *b*, we compute the cell strain *ε* using Eq. 10 (Fig. 2a). We also compute the local fluid shear stress *σ* (*y*_0_) for a cell-free fluid at the cell’s centroid position using Eq. 4 (Fig. 1f).

**Figure 1.**
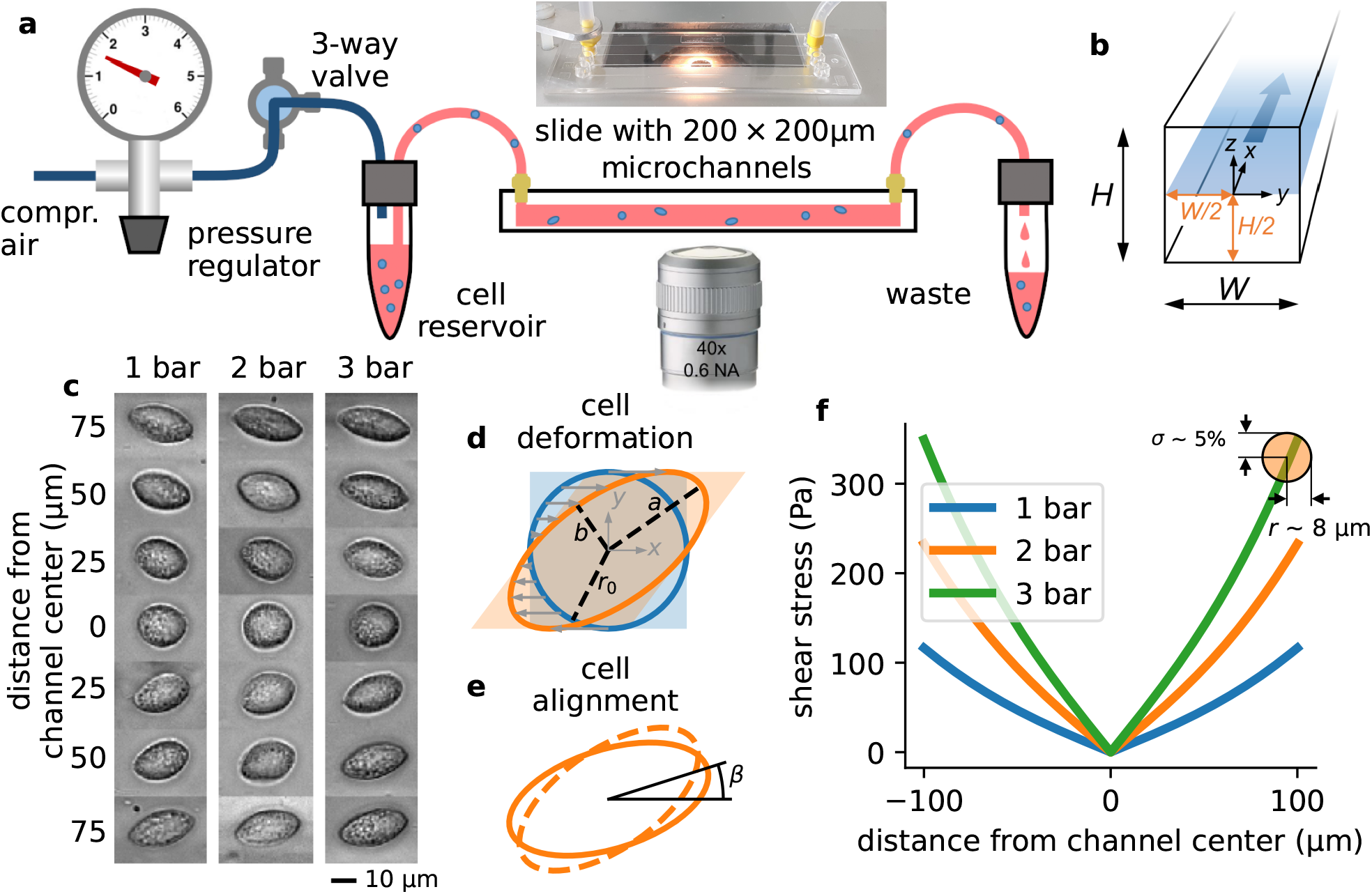
Measurement setup and principle. **a**, Schematic of the microfluidic device. **b**, Cross section through the microchannel with dimensions *W* = *H* = 200 µm. The focal plane of the microscope at a height of *H*/2 = 100 µm is indicated by the blue shaded area. Fluid flow is in *x* direction. **c**, Bright field images of NIH-3T3 cells under control conditions at different y-positions in a microchannel at a pressure of 1, 2, and 3 bar. Cells appear round in the channel center and become more elongated near the walls. **d**, Illustration of cell deformations under fluid shear. The circular cell with radius *r*_0_ (blue) is transformed to an elliptical shape (orange) with semi-major axis *a* and semi-minor axis *b* depending on the ratio of fluid shear stress and the cell’s shear modulus (Eq. 16). **e**, The sheared cell (dashed outline) will partially align in flow direction (solid outline), characterized by an alignment angle *β*. This angle depends on the ratio of cell viscosity and suspension fluid viscosity (Eq. 17). *a, b*, and *β* are measured from the segmented cell shapes. **f**, Fluid shear stress (computed according to Eq. 4) versus distance from the channel center in *y*-direction for three different pressures of 1, 2 and 3 bar. The shear stress varies by maximally 5% across the cell surface for a typical cell with a radius of 8 µm (indicated by the orange circle).

**Figure 2.**
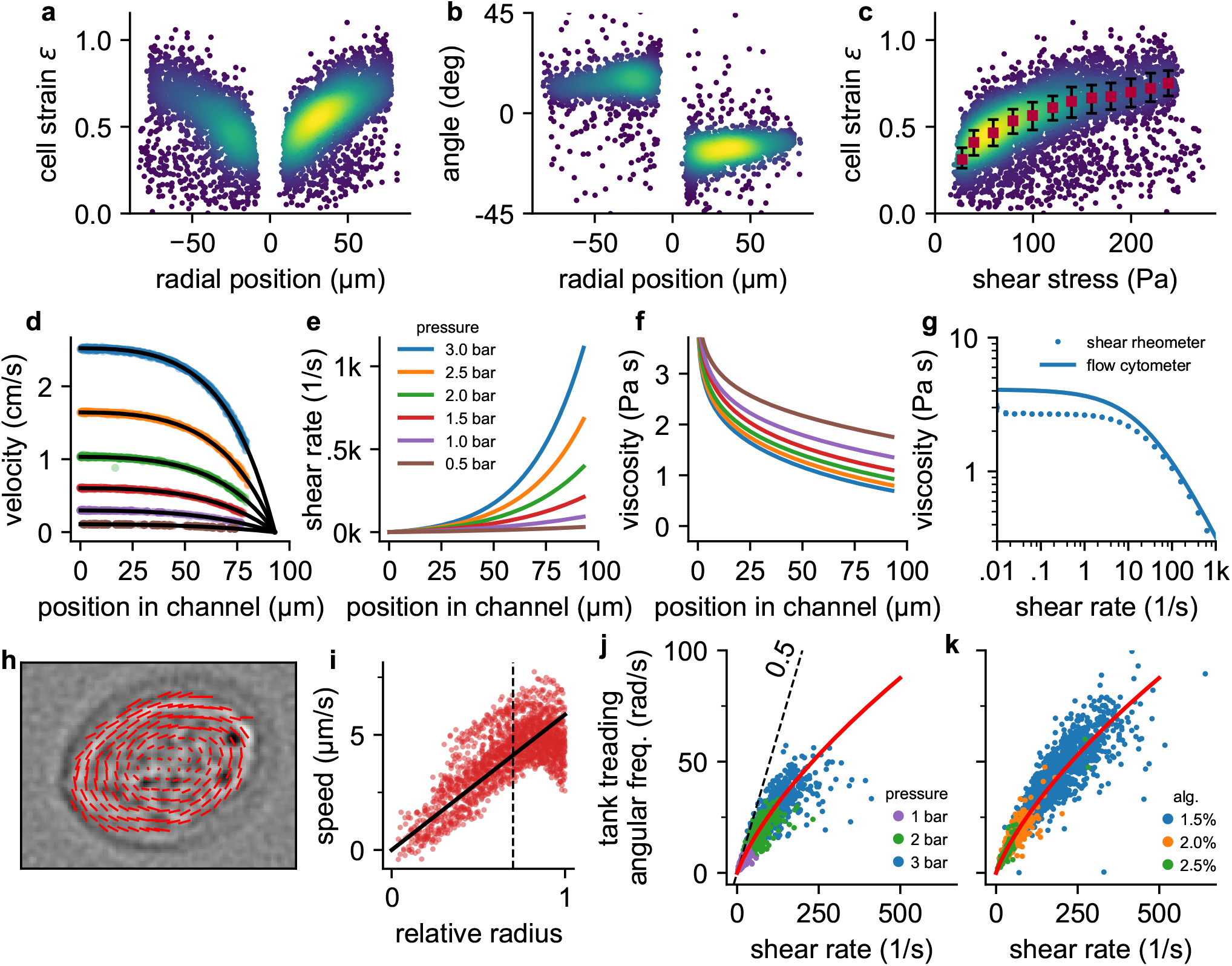
Cell responses to shear stress and shear rate. **a**, Cell strain versus radial (*y*) position in the channel for NIH-3T3 cells at a pressure of 3 bar. Each data point corresponds to a single cell. Colors indicate Gaussian kernel density. **b**, Cell alignment angle *β* versus radial position in the channel (*y*) for the same cells as in **a. c**, Cell strain versus shear stress for the same cells as in **a**. Red squares indicate median values over shear stress bins of 20 Pa starting from 10 Pa, error bars indicate quartiles. **d**, Fluid flow velocity versus radial channel position (*y*) for different driving pressures (0.5, 1.0, 1.5, 2.0, 2.5, 3.0 bar). Each data point corresponds to the speed of a single cell. Black lines show individual fit curves obtained by fitting the Cross-model (power-law shear thinning fluid with zero-stress viscosity) to the velocity profile (Eq. 5 - Eq. 9). **e**, Shear rate of the suspension fluid versus radial channel position (*y*) for different driving pressures. The shear rate is computed with Eq. 7. **f**, Local suspension fluid viscosity at different channel positions computed with Eq. 6. **g**, Suspension fluid viscosity versus shear rate from the fit of the Cross-model (blue line) to the data shown in **d**, and measured with a cone-plate rheometer (blue circles). **h**, Tank-treading rotation of a cell in the channel, quantified from the optical flow between two subsequent images. **i**, Rotational speed of cell image pixels (same cell as in **h**) versus the ellipse-corrected radius (radial pixel position normalized by the radius of the cell ellipse at that angle). Only cell pixels with an ellipse-corrected radius below 0.7 (dotted line) are used for the linear fit of the tank-treading frequency to the data (solid line) to avoid cell boundary artefacts. **j**, The angular tank-treading frequency *ω*_tt_ increases with the shear rate, with a slope approaching 0.5 for small shear rates (dashed black line). Each point represents the data of an individual cell; different colors indicate different pressures. The red line presents the fit of Eq. 20 to the data. **k**, same as in **j** but for measurements at a pressure of 2 bar in differently concentrated alginate hydrogels.

#### Cell deformations under fluid shear stress

Cells are nearly circular in the center, and they elongate and align in flow direction near the channel walls (Fig. 1c, 2a,b) where they are exposed to higher fluid shear stress (Fig. 1f). Cells imaged at the same position within the channel also tend to become more elongated with increasing pressure (Fig. 1c). When we plot cell strain *ε* versus shear stress *σ* across the microfluidic channel (Fig. 2c), we find that the cell strain increases non-linearly with increasing fluid shear stress. In particular, the slope of the strain versus stress relationship decreases for higher stress values. This behavior is predominantly due to a dissipative process caused by the tank tread-like motion of the cells.

#### Tank-treading

The radial velocity gradient of the flow field (the shear rate 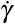) creates a torque on the sheared and elongated cells and causes them to align in flow direction (Fig. 1e, 2b) and to rotate in a tank-treading manner (SI video 1): the cell’s elongated shape and alignment angle *β* remain stationary, but internally, the cell is constantly rotating as if being kneaded between two plates^4^.

From a series of images that show the same cells as they flow through the channel, we compute the radial velocity profile *v*(*y*) of the fluid flow (Eq. 9, Fig. 2d), the shear rate profile 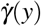 (Eq. 7, Fig. 2e), and the tank-treading frequency *ω*_tt_ of each cell (Fig. 2h,i). We find that the tank-treading frequency of a cell is zero at the channel center and increases towards the channel walls (Fig. 2j,k). At low shear rates (low driving pressure or near the channel center), the rotation rate *ω*_tt_/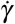 of individual cells is close to the Einstein-limit of 1/2, as theoretically predicted for spheres that are tank-treading in a Newtonian fluid^1,5^. Tank-treading dissipates energy in proportion to the cell’s internal viscosity, rotation frequency, and strain. This energy dissipation therefore limits the cell strain in regions of high shear rate and hence shear stress (Fig. 2c).

### Viscoelastic model

We can quantitatively explain the non-linear strain-stress relationship (Fig. 2c) and its pressure-dependency by a theoretical framework describing the deformation and alignment of viscoelastic spheres in a viscous fluid under steady shear flow^1^. This theoretical framework (in the following referred to as Roscoe-theory) predicts that the cell strain *ε* increases proportional with the shear stress *σ* and the sine of the alignment angle *β*, and inversely proportional with the elastic modulus *G*^′^ of the cell (Eq. 16). The alignment angle *β* in turn depends on the cell’s loss modulus *G*^′′^, the local shear rate 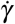 and the local viscosity *η* of the suspension fluid (Eq. 17). With increasing elastic modulus, cells are predicted to deform less (smaller strain *ε*) and to align less in flow direction (larger alignment angle *β*) when exposed to a fixed shear stress and shear rate. With increasing loss modulus, cells are also predicted to deform less but to align more in flow direction. Thus, from the measurements of cell strain, alignment angle, local shear stress, local shear rate, and local viscosity, Roscoe-theory allows us to compute the viscoelastic properties (*G*^′^(*ω*) and *G*^′′^(*ω*)) of individual cells at their specific angular tank-treading frequency.

### Power-law behavior of cells

When we plot *G*^′^ and *G*^′′^ of individual cells versus their tank-treading frequency *ω*_tt_ (Fig. 3a), we find that the complex shear moduli 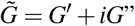 of a cell population approximately follow a power-law relationship of the form

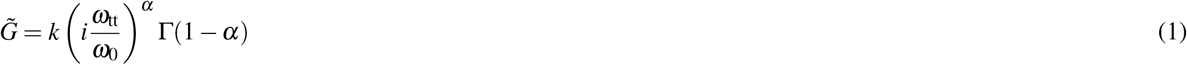

where Γ is the Gamma-function, *k* is the elastic shear modulus (cell stiffness) referenced to an arbitrarily chosen frequency *ω*_0_ (here we chose 1 rad/s), *α* is the power-law exponent that characterizes the fluidity of the cell (zero indicating purely Hookean elastic behavior, unity indicating Newtonian viscous behavior), and 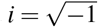^6^. Such a behavior of a cell population emerges if the rheology of individual cells also follows a power-law relationship. Thus, using Eq. 1, we can compare the mechanical behavior of cells measured at different tank treading frequencies by computing their stiffness *k* (using Eq. 21) and fluidity *α* (using Eq. 22).

**Figure 3.**
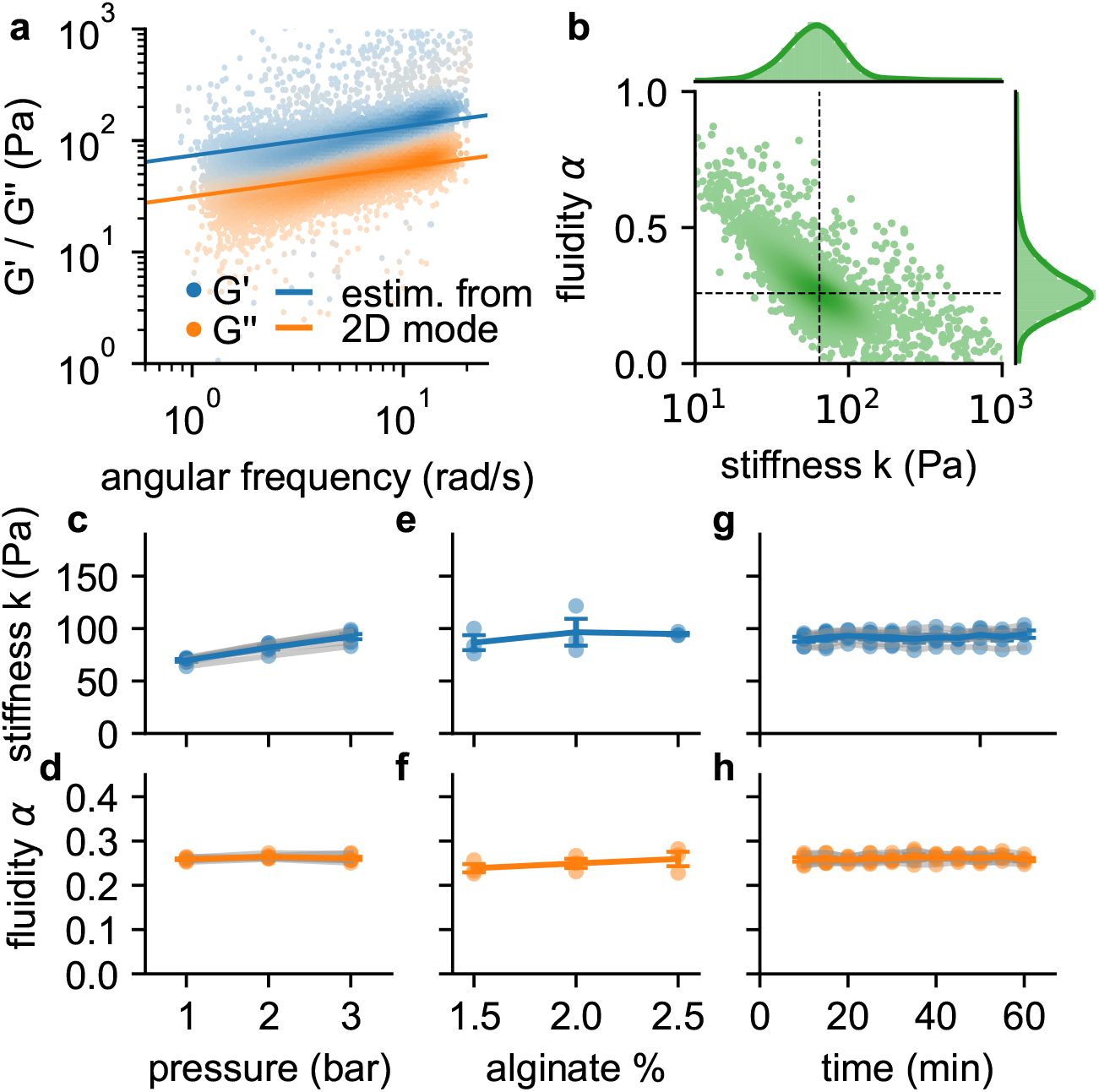
Frequency, pressure, suspension fluid and time dependency of viscoelastic cell behavior. **a**, *G*^′^ (blue dots) and *G*^′′^ (orange dots) of individual NIH-3T3 cells measured at 300 kPa. Lines are not a fit to the data but indicate the predicted behavior of *G*^′^ (blue line) and *G*^′′^ (orange line) versus angular (tank-treading) frequency according to Eq. 1 of a typical cell with stiffness and fluidity corresponding to the mode of the 2D histogram shown in **b. b**, Distribution of stiffness *k* and fluidity *α* of the same cells as shown in **a**, dashed lines indicate the mode of the 2D histogram. Color coding shows 2D Gaussian kernel density estimation. Histograms show the probability density distributions of *k* (top) and *α* (side) with Gaussian kernel density estimates (green line). **c**, Stiffness *k* of NIH-3T3 cells increases with pressure (blue lines and symbols indicate mean±se, gray lines and transparent symbols indicate individual data from 6 independent measurements). **d**, Fluidity *α* (same cells as in **c**) remains constant for all measured pressures. **e**,**f**, Stiffness and fluidity show only a weak dependence on alginate concentration (measured at a pressure of 200 kPa, mean±se (blue) from 3 independent measurements (gray)). **g**,**h**, *k* and *α* of NIH-3T3 cells remain constant for at least 60 min after suspending them in a 2% alginate solution (measured at a pressure of 300 kPa, mean±se (blue) from 5 independent measurements (gray)).

We find in agreement with previous reports^7–11^ that the individual stiffness values *k* are typically log-normal distributed, and the fluidity values *α* are normal distributed (Fig. 3b). Moreover, also in agreement with previous reports, we find an inverse relationship between stiffness and fluidity, whereby stiffer cells tend to be less fluid-like^6,12,13^. Due to this coupling, the mode of the two-dimensional distribution of *α* and *k* (the most common combination of *α* and *k* among all cells, as estimated from the maximum of the Gaussian kernel-density, Fig. 3b), provides a robust measure for the mechanical behavior of a cell population.

The slope of the *G*^′^ and *G*^′′^ versus frequency relationship on a logarithmic scale (Fig. 3a), which is the power-law exponent *α* of the cell population, appears larger than the power-law exponent *α* of individual cells computed with Eq. 22 (solid lines in Fig. 3a). A likely explanation for this behavior is a shear stress-induced stiffening of the cells: Since regions in the channel with higher shear stress also have a higher shear rate and hence induce a higher tank-treading frequency, *G*^′^ and *G*^′′^ of stress-stiffening cells seemingly increase faster with frequency than predicted by Eq. 1.

### Stress stiffening

To test if suspended NIH-3T3 cells exhibit stress stiffening, we increase the driving pressure from 100 kPa to 300 kPa, which increases the maximum shear stress at the channel wall from 116 Pa to 349 Pa (Fig. 1f). Cell fluidity remains constant over this pressure range, but the median stiffness of the cell population increases with increasing pressure by 33% (Fig. 3c,d). To explore to which extent this stiffness increase is caused by a higher shear stress as opposed to a higher shear rate, we keep the pressure constant at 200 kPa but increase the alginate concentration from 1.5% to 2.5% and therefore the viscosity of the suspension medium from 2.2 Pa·s to 9.2 Pa·s (zero-shear viscosity *η*_0_ as determined with Eq. 6). This causes the shear rate to decrease and leads to a slight but not statistically significant increase in stiffness and fluidity (Fig. 3e,f). Hence, the increase of cell stiffness at a higher driving pressure is induced by stress-stiffening and not by a higher shear rate. We also verify that cell stiffness and fluidity remain stable over a period of up to 60 min after suspending the cells in a 2% alginate solution (Fig. 3g,h).

### Validation with polyacrylamide beads

To evaluate the accuracy of our method, we measure 16 µm diameter polyacrylamide (PAAm) beads with three different nominal stiffnesses, in a range similar to living cells (Fig. 4a-c). The frequency-dependency of *G*^′^ and *G*^′′^ of the beads are calibrated using oscillatory atomic force microscopy (AFM), and conform to the relationship

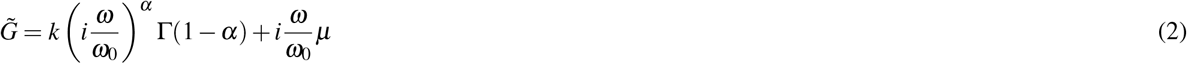

with *k* = 120 Pa and *α* = 0.041 (Fig. 4d). Using shear flow cytometry, we obtain similar values (*k* = 141 Pa and *α* = 0.09). Moreover, *k* and *α* are found to be largely pressure-independent (from 0.2–2 bar) (Fig. 4e), as expected for a linear material such as PAAm. By lowering the acrylamide-bisacrylamide total monomer concentration (*C*_AAmBis_) we produce softer beads with stiffness and fluidity values of of *k* = 22 Pa, *α* = 0.11 for 5.9% *C*_AAmBis_, and *k* = 5.4 Pa, *α* = 0.12 for 3.9% *C*_AAmBis_, as measured by AFM indentation. Stiffness and fluidity measurements using shear flow deformation cytometry also give comparable values, with *k* = 17 Pa for 5.9% *C*_AAmBis_, and *k* = 11 Pa for 3.9% *C*_AAmBis_ (SI Fig. S3). Fluidity is close to zero for strains below unity (*α* = 0.0018 for 5.9% *C*_AAmBis_, and *α* = 0.039 for 3.9% *C*_AAmBis_), indicating predominantly elastic behavior as expected, but fluidity increases at higher strains (Fig. 4f) due to fluid-induced (poroelastic) relaxation processes^14^. Together, these results demonstrate that our method provides quantitatively accurate estimates for the elastic and dissipative properties of soft spherical particles.

**Figure 4.**
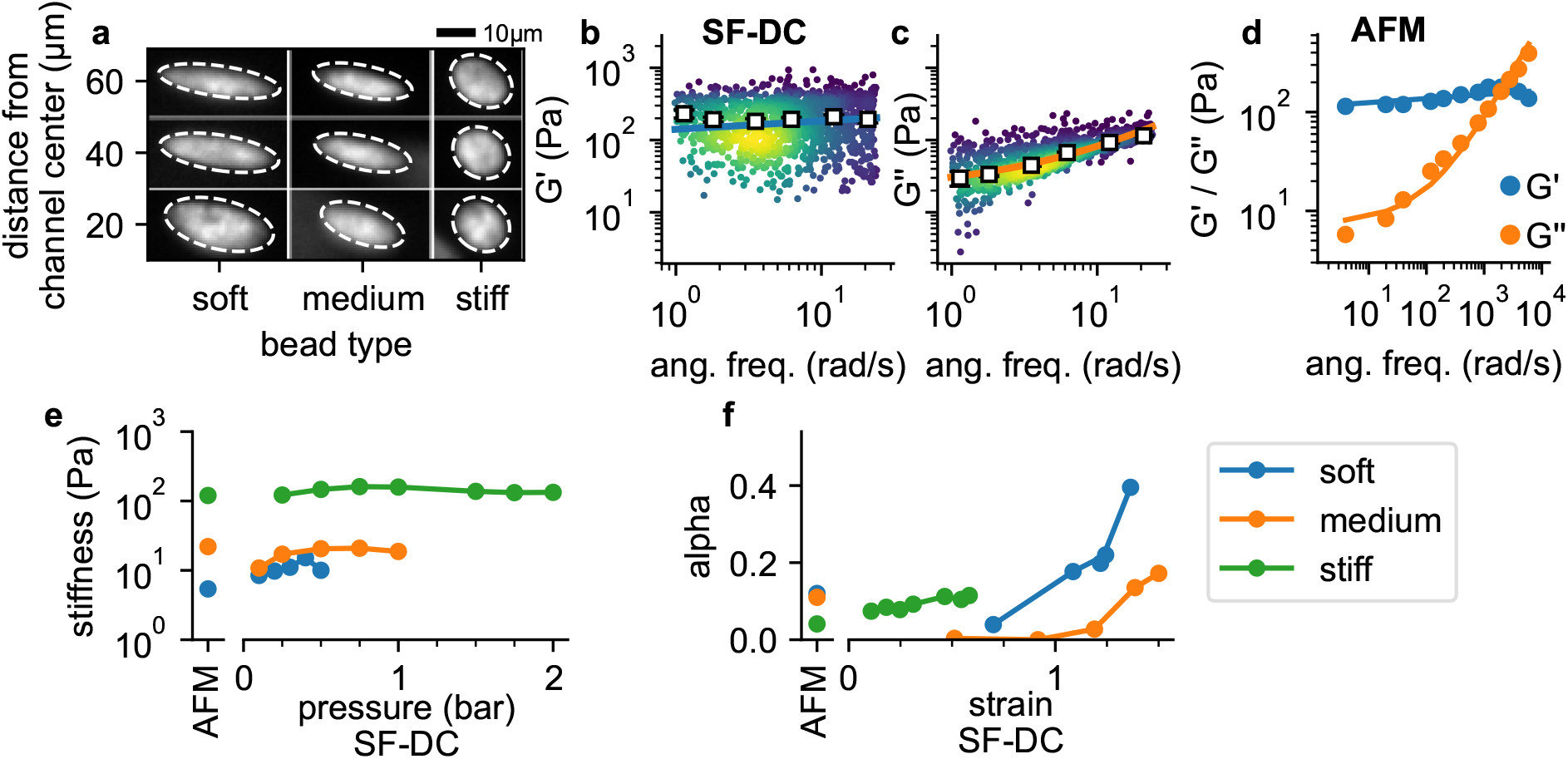
Validation with polyacrylamide beads. **a**, Deformation of PAAm beads with different acralymide-bisacrylamide total monomer concentrations (soft 3.9%, medium 5.9%, stiff 6.9%) at different positions in the channel. *G*^′^ (**b**) and *G*^′′^ (**c**) for stiff beads at 2 bar. White squares indicate binned median values, blue and orange solid lines are the fit of Eq. 2 to the data. **d**, AFM data (*G*^′^ and *G*^′′^ versus frequency, mean values from 14 stiff PAAm beads (blue/orange circles), solid lines are the fit of Eq. 2 to the data). **e**, AFM-measured stiffness compared to the stiffness versus pressure measured with shear flow deformation cytometry (SF-DC) for differently stiff PAAm beads. **f**, AFM-measured fluidity compared to fluidity versus strain measured with SF-DC for the same beads as in **e**.

We next compare the viscoelastic properties of monocytic THP-1 cells probed by shear flow cytometry and atomic force microscopy (AFM). We acquire force-indentation curves at rates of ∼1/s (Fig. 5c), which is within the range of strain rates that cells experience in our shear flow cytometry setup. AFM measurements show that THP-1 cells conform to power-law rheology according to Eq. 2, from which we extract the shear modulus *k* and fluidity *α* (Fig. 5b). The shear modulus values of THP-1 cells measured with shear flow cytometry (*k* = 87 Pa, *α* = 0.19) are in reasonable agreement with those from AFM measurements (*k* = 120 Pa, *α* = 0.21).

**Figure 5.**
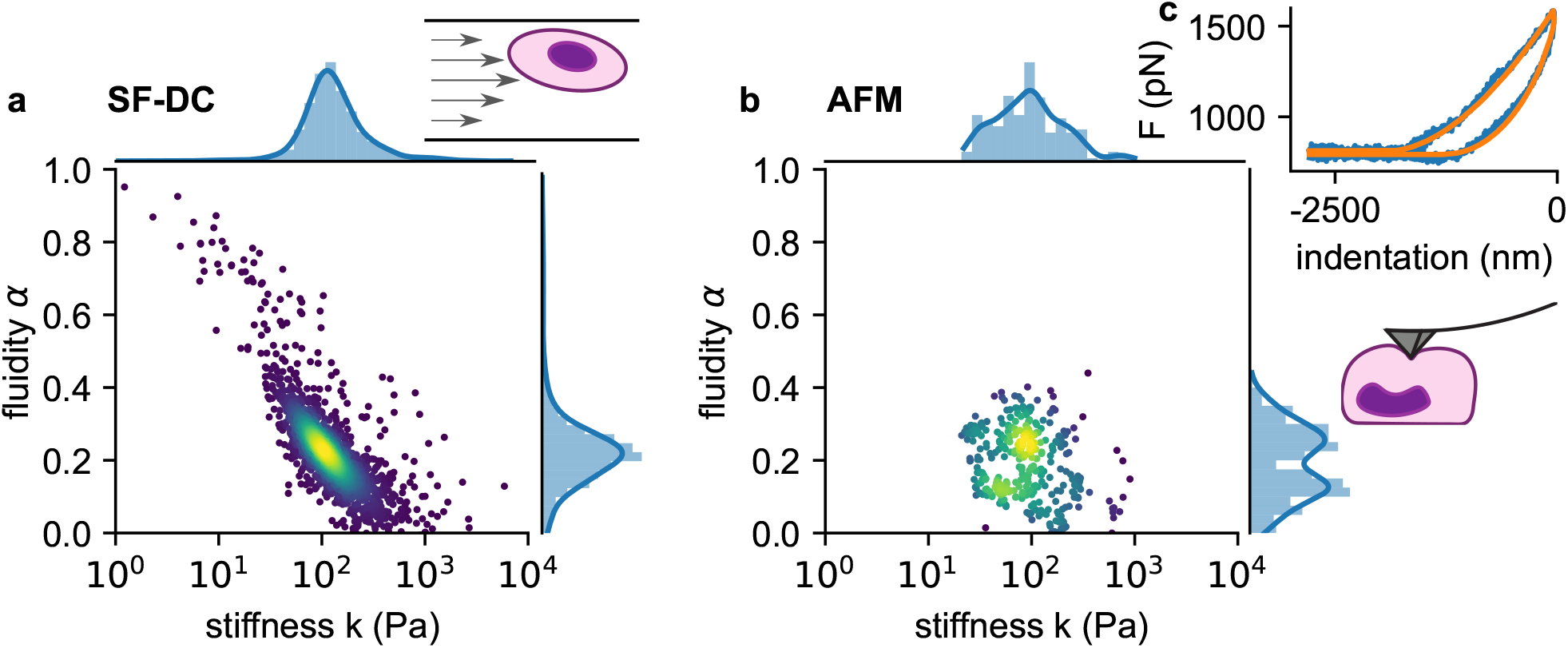
Comparison of viscoelastic cell properties measured with shear flow deformability cytometry (SF-DC) and AFM. **a**. Stiffness *k* versus fluidity *α* of THP1 cells (n=5000) measured with SF-DC at a pressure of 2 bar, 2% alginate solution. Colors represent Gaussian kernel density. Histograms show the probability density distributions of *k* (top) and *α* (side) with Gaussian kernel density estimates (blue line). **b**, AFM measurements of THP1 cells. Each point represents *k* and *α* from one cell, each obtained from the fit of Eq. 24 to 3 or more force-indentation curves for each cell. **c**, Typical force-indentation curve (blue line) and fit with Eq. 24 (orange line).

### Dose response measurements

We perform dose-response measurements using latrunculin B (LatB), which prevents the polymerization of monomeric actin and leads to a depolymerization of the actin cytoskeleton^2^. NIH-3T3 fibroblasts soften with increasing doses of LatB (1-1000 nM) according to a sigmoidal (Hill-Langmuir) relationship, with a maximum response of 1.56-fold and a half-maximum dose of EC50 = 31.2 nM. These responses agree with published data obtained using real-time deformability cytometry (RT-DC) measurements on HL-60 cells (maximum response 1.46-fold, EC50 = 26.5 nM)^2^. When we measure pro-myoblast HL-60 suspension cells with our setup, EC50 is similar to published data (25.9 nM), but the maximum response is much higher (5.4 fold).

### Role of intermediate filaments

To explore the attenuated LatB responsiveness of NIH-3T3 fibroblasts compared to HL-60 leukemia cells, we reasoned that NIH-3T3 cells express high levels of the intermediate filament protein vimentin (Fig. 7a) that may protect the cells from excessive deformations when filamentous actin is depolymerized. To test this idea, we measure the stiffness of NIH-3T3 and vimentin-knock-out (vim(-/-)) mouse embryonic fibroblasts (MEFs) in response to 30 min treatment with cytochalasin D (2 µM), which binds to the barbed end of filamentous actin and—similar to LatB—leads to a net depolymerization of the actin cytoskeleton (Fig. 7a). We find that cytochalasin D treated vim(-/-) cells soften by a considerably greater extent (2.16-fold) compared to wild-type cells (1.22 fold) (Fig. 7b,c), in support of the notion that vimentin stabilizes the cytoskeleton.

**Figure 6.**
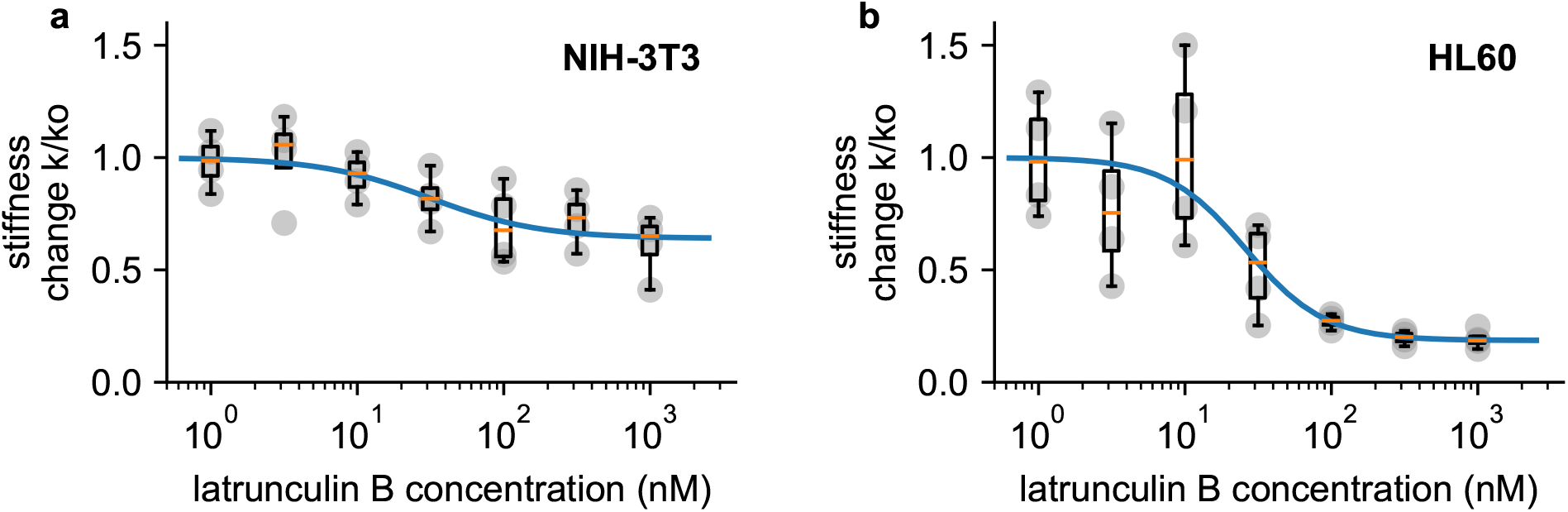
Dose response measurements. **a**, Dose response curve of NIH-3T3 cells treated with different concentrations of latrunculin B. Stiffness is normalized to the stiffness of DMSO-treated cells, grey points indicate n = 4 independent measurements for each concentration, each measurement is the average of a 0.5, 1, 2, and 3 bar measurement, boxplot indicate median (orange line) and 25 and 75 percentiles, whiskers indicate 5 and 95 percentiles. Blue line is the fit of the Hill-Langmuir-equation to the data, with an EC50 of 31.2 nM. **b**, Dose response curve of HL 60 cells. Hill-Langmuir fit gives an EC50 value of 25.9 nM.

**Figure 7.**
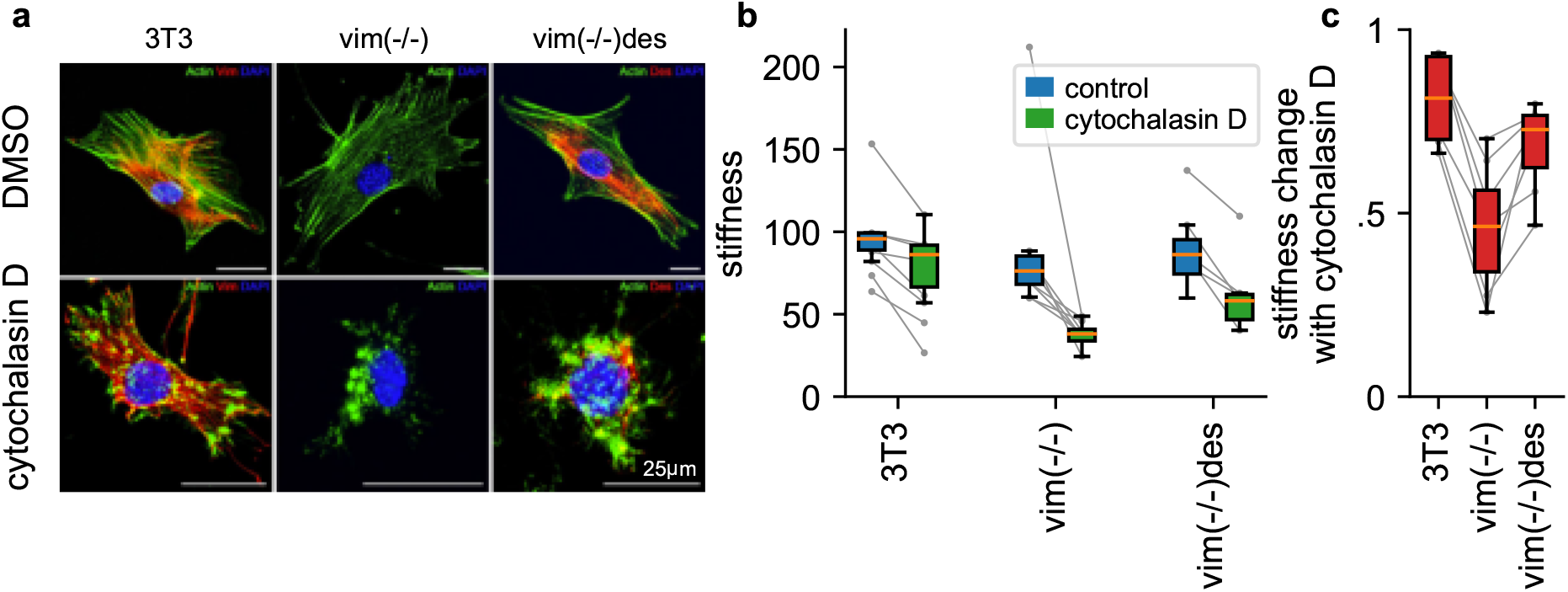
Influence of intermediate filaments. **a**, NIH-3T3, vimentin-knockout, and desmin-knockin MEFs in DMSO control conditions (upper row) and with cytochalasin D treatment (lower row). Actin is shown in green, desmin in red and the nucleus in blue. **b**, Stiffness of DMSO control (blue box) and cytochalasin D treated cells (green box) (orange line: median, box: 25 and 75 percentile, whiskers: 5 and 95 percentile, gray points and lines connect mean values from independent measurements performed on the same day). **c**, Stiffness change after treatment with cytochalasin D relative to DMSO control.

To explore if the cytoskeleton-stabilizing effect of vimentin is a general feature also of other intermediate filament networks, we measure the cytochalsin-D response of desmin-transfected embryonic fibroblasts from vimentin knock-out mice (vim(-/-)des). Desmin, which is the dominant intermediate filament in skeletal muscle, forms an intermediate filament network in fibroblasts that is structurally similar to the vimentin network in wild-type cells (Fig. 7a). Similar to wild-type cells, vim(-/-) desmin-expressing MEFs also display an attenuated cytochalasin D response (1.37 fold), confirming that both the vimentin and desmin intermediate filament network can protect cells from excessive deformations when filamentous actin is depolymerized.

### Cell Cycle dependence

In our measurements, we observe that larger NIH-3T3 cells tend to be softer compared to smaller cells (Fig. 8a). We hypothesized that this weak size-dependence of cell stiffness might be attributable to cell cycle progression, which leads to changes in chromatin compaction and cell volume. To test this hypothesis, we extend our setup to acquire green fluorescent images alongside bright field images of cells transfected with a two-color fluorescent Fucci cell cycle indicator^15^. Fucci-transfected cells display high red and low green fluorescence when they are in G1 phase, and low red but increasing levels of green fluorescence as they progress into S, G2 and early M-phase^15^. We measure the cell cycle distribution of NIH-T3T cells before harvesting using epifluorescence microscopy (Fig. 8b), and map the distribution to the green fluorescent intensities measured in our shear flow cytometry setup (Fig. 8c).

**Figure 8.**
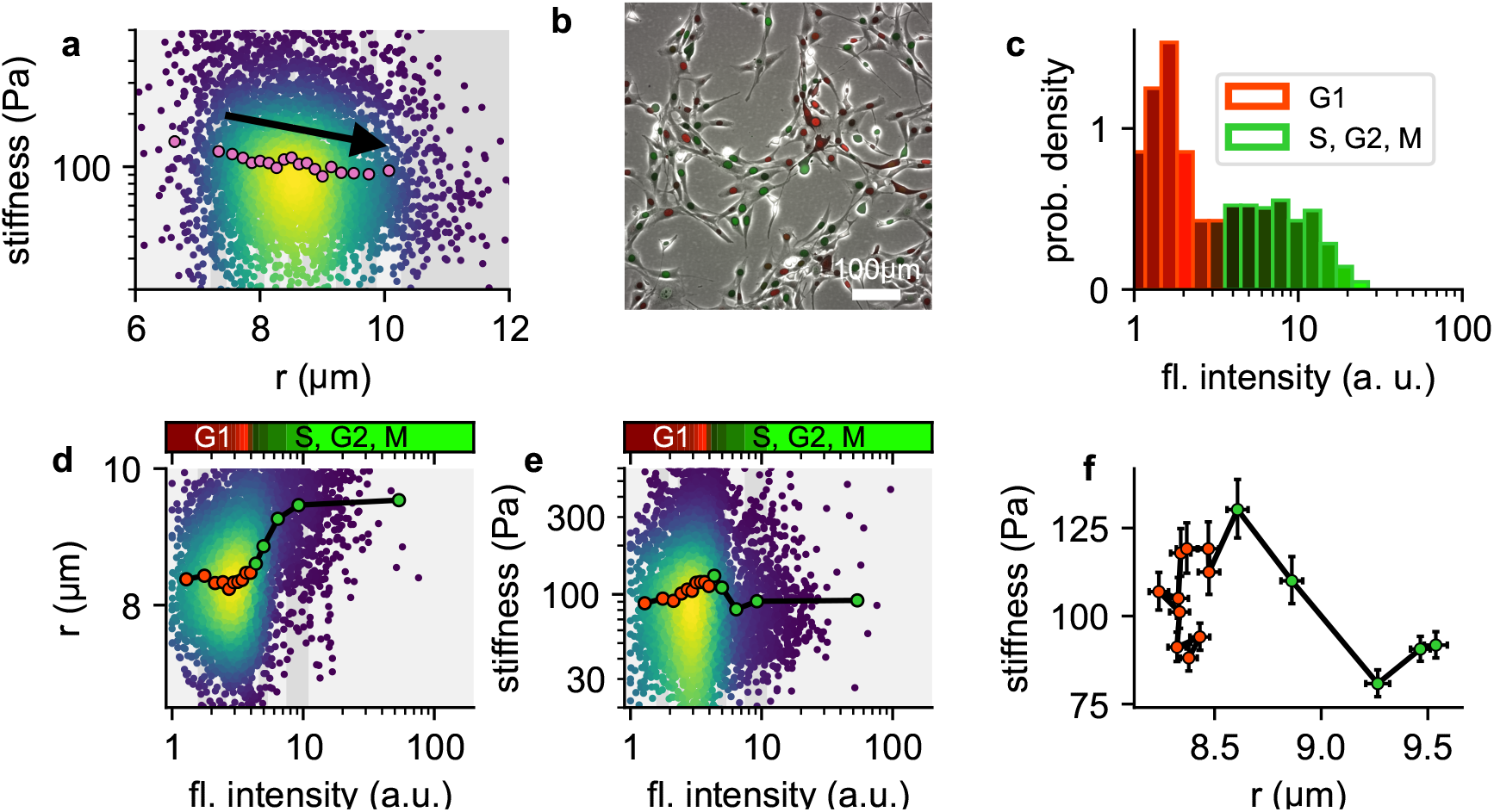
Influence of cell cycle. **a**, Cell stiffness versus cell radius, each point corresponds to data from one cell, colors represent Gaussian kernel density, pink circles show median values over bins with equal cell count. Cell stiffness tends to decrease with increasing cell radius. **b**, Phase contrast and fluorescent image of Fucci-cell cycle indicator-transfected NIH-3T3 cells. Cells in G1 phase show low green and high red fluorescence intensities, cells in S, G2 or M phase show high green and low red fluorescence intensities. **c**, Histogram of green fluorescence intensities of Fucci-cell transfected NIH-3T3 cells. Bar colors reflect the RGB-colormap of the red and green channel intensities averaged over all cells within a bin. Accordingly, the cell cycle can be deduced from the green intensity alone. **d**, Cell radius versus green fluorescent intensity. Each point corresponds to data from one cell, colors represent Gaussian kernel density, circles show median values over bins containing an equal number (∼ 100) of cells. Colorbar represents the RGB-colormap of the red and green intensities of the cells before harvesting, mapped onto the green fluorescent intensity after harvesting measured in the shear flow cytometer. Cell radius increases after cells exit G1 phase. **e**, Cell stiffness versus green fluorescent intensity. Cells stiffness increases during G1 phase and decreases after entering S phase. **f**, Cell stiffness versus cell radius; data points correspond the the median values in **d** and **e**, red color designates cells in G1 phase, green color designates cells in S, G2 or early M phase. During G phase, cells increase their stiffness while maintaining their radius. After entering S phase, cells increase their radius while their stiffness decreases.

We find as expected that cell radius increases with cell cycle progression (Fig. 8d). In addition, cell stiffness steadily increases towards the end of the G1 and the beginning of the S-phase, and then rapidly decreases as the cell cycle progresses (Fig. 8e). When we bin the cells according to their green fluorescent intensities (i.e. according to their cell cycle progression) and plot stiffness versus cell radius (Fig. 8f), we find substantially larger and non-monotonic fluctuations of cell stiffness versus cell radius, compared to the smaller, monotonic decrease of cell stiffness in the radius-binned data (Fig. 8c). These differences arise because changes in cell stiffness and cell radius occur at different stages of the cell cycle.

## Discussion

Viscoelastic cell properties can be measured with established methods such as atomic force microscopy^16^, micropipette aspirations^17^, or magnetic tweezer microrheology^11^. These methods have a relatively low throughput of typically below 10–100 cells/h. The need to measure cell mechanical properties with substantially higher throughput led to the recent development of various microfluidic techniques^2^ including hydrodynamic stretching^18^, real-time deformability cytometry^19,20^, micro-filtration^21^, and micro-constriction systems^13,22^.

Our method builds on previously established high-throughput microfluidic approaches, with several modifications: We suspend cells in a medium that is pumped with high pressure (typically 50–300 kPa) through a commercially available long, parallel microfluidic channel with one inlet and outlet (no flow-focussing geometry is needed). This large driving pressure gives rise to sufficiently large (> 50 Pa) shear stresses to induce measurable cell deformations. The high pressure can be controlled with a simple pressure regulator, without the need for a precise microfluidic controller. The width and height of the channel (200 µm) are much larger than the cell diameter, which prevents clogging due to debris and ensures that fluid shear stresses do not vary appreciably across the cell diameter. By suspending the cells in a fluid with high viscosity (typically > 1 Pa·s), we achieve a flow speed that is sufficiently low (< 20 mm/s) so that the cells’ speed, position, and shape can be captured without motion blur at a typical exposure time of 30 µs using a standard CMOS-camera mounted to a routine laboratory microscope.

The lateral flow profile in the channel causes a tank-treading-like cell motion, which imposes periodic cell deformations with frequencies on the order of 10 Hz. At such low frequencies and strain rates, elastic cell properties dominate over viscous cell properties^6,23^. The cell transit through the microfluidic channel lasts for several seconds, which is much longer than the period time of the cells’ tank treading rotation, implying that the measured cell deformations can safely be assumed to have reached a steady-state.

From images of the same cell as it is flowing through the channel, we estimate the tank treading frequency and the flow velocity; from the flow velocity profile across the channel, we compute the local shear rate (Eq. 5) and the local viscosity of the suspension fluid (Eq. 6); from the radial cell position, we compute the local shear stress (Eq. 4); from the cell shape, we compute the strain (Eq. 10) and the alignment angle in flow direction. From these measurements, we finally compute the cell’s viscoelastic properties (stiffness and fluidity, Eq. 21 and 22). Hence, once the flow velocity profile is known, we can determine a cell’s viscoelastic properties from a single image, which is the main difference compared to alternative microfluidic techniques for measuring viscoelastic cell properties, where cell shape changes in response to a sudden increase or decrease in shear stress are recorded and analyzed^13,20^.

Our computation is based on a theoretical model proposed by R. Roscoe that describes the deformation of homogeneous, isotropic, incompressible neo-Hookean viscoelastic spherical particles under fluid shear stress^1^. Cells in suspensions, however, are known to deform non-linearly^22^. They do not consist of a homogeneous material but of different components (e.g. the cell cortex and the nucleus) with different mechanical properties^16,17,21,23^. Moreover, cells do not always deform into ellipsoidal shapes but occasionally deform into sigmoidal shapes, which becomes more pronounced in response to larger shear stresses or drugs that soften the cytoskeleton, such as cytochalasin D or latrunculin B.

Despite the simplified assumptions of the Roscoe theory, however, our cell rheological measurements agree with previously published findings that were obtained using a range of different methods and models, namely that suspended cells show a behavior that is consistent with power-law rheology, that the elasticity of individual cells is log-normal distributed, that the fluidity of individual cells is normal-distributed, and that stiffness and fluidity scale inversely^6,7,13,24^. These experimental findings are in agreement with predictions from soft glassy rheology^6,25^. Moreover, we show that stiffness and fluidity of polyacrylamide beads and cells measured with shear flow deformation cytometry agree quantitatively with AFM measurements.

Our measurements are insensitive to changes in the viscosity of the suspension medium, demonstrating that the fluid-mechanical assumptions of the Roscoe theory hold in the case of living cells in a shear-thinning suspension fluid. We find that cells appear stiffer when measured at higher driving pressures, likely due to stress- or strain-stiffening of the cells^22^, which is ignored in the Roscoe theory. When we measure linearly elastic polyacrylamide beads over a 10-fold pressure range (from 20–200 kPa), we see a constant, pressure-independent shear modulus and agreement with the stiffness and fluidity values measured using AFM, demonstrating that the Roscoe theory gives quantitatively accurate estimates, regardless of driving pressure and suspension fluid viscosity.

Roscoe theory estimates the cell viscosity relative to the viscosity of the suspension fluid, which for a shear thinning fluid such as alginate can be difficult to measure. However, since we know the fluid profile in the microfluidic channel (from the flow speed of hundreds of cells), we can—as a by-product of our method—estimate the rheological properties of the suspension fluid, including its shear thinning behavior. The rheological parameters for alginate solutions measured this way closely agree with cone-plate rheometer measurements, with relative deviations of 31% over a shear rate spanning 5 orders in magnitude (from 0.01–1000 s^−1^).

Our method measures each cell at a single tank-treading frequency that depends on the cell’s lateral position in the channel. Thus, we sample the frequency-dependent mechanical properties of a cell population by observing many cells at different channel positions. The tank-treading frequency can be directly measured using particle flow analysis methods in a subset of the cells that shows small features with high contrast. For the remaining cells, it is possible to estimate the tank treading frequency from the local shear rate according to an empirical equation (Eq. 20). This equation holds for the cell types and suspension fluids used in our study, but we do not claim that it holds universally for other cell types and suspension fluids.

To demonstrate its practical applicability, we apply our method to measure the stiffness of HL-60 cells in response to different doses of the actin-depolymerizing agent latrunculin B. We find in agreement with previous observations a half-maximum dose (EC50) of around 30 nM, but a considerably larger softening of the cells by a factor of 5.4 fold at the highest dose of 1 µM, compared to a softening of only 1.5 fold that is seen with other microfluidic techniques (constriction microfluidic constriction-based deformability cytometry (cDC), and real-time deformability cytometry (RT-DC)^2^. This higher responsiveness is likely attributable to the relatively low cellular strain rate of our method, which is on the order of 10 Hz, compared to strain rates of around 100 Hz in the case of RT-DC. At these high strain rates, viscous cell behavior starts to dominate over cytoskeleton-associated elastic responses^6,23^. Accordingly, when cells are measured with extensional flow deformability, a method that operates at even higher strain rates in the kHz-range, they do not appreciably soften in response to LatB^2,18^.

We also demonstrate that the cell softening induced by cytochalasin D, another actin-depolymerizing drug, is attenuated in the presence of intermediate filaments (vimentin or desmin), and becomes more pronounced when intermediate filaments are absent. This finding is in support of the notion that intermediate filaments protect mechanically stressed cells against excessive strain and damage^26^.

Shear stress deformability cytometry can be combined with fluorescent imaging. Here, we image the viscoelastic properties of NIH-3T3 cells together with the cell cycle using the fluorescent Fucci indicator. Our data demonstrate that NH-3T3 cells stiffen during the course of cell cycle progression in G1 phase, with a maximum stiffness during late G1 – early S-phase, and then soften before they enter the G2 and M-Phase. Since cell volume also increases during the transition from G1 to S phase, we find a slight overall dependence of cell stiffness on cell size in the case of NIH-3T3 cells (Fig. 8c). This cell size dependence is also detectable in HL-60 and THP1-cells (SI Fig. S2).

In summary, shear flow deformation cytometry provides accurate quantitative measurements of viscoelastic cell properties at high throughput. The method can be easily and inexpensively implemented on standard or research grade microscopes. We provide user-friendly software for image acquisition and data analysis, which can be downloaded at https://github.com/fabrylab/shear_flow_deformation_cytometer. Currently, the method stores the acquired uncompressed images on a hard drive, which in the case of typically 10,000 images for a single experiment lasting 20 s amounts to a storage space of nearly 4 GB. The image data are analyzed afterwards, which at a rate of around 50 images per second can take several minutes. Future software developments and faster computer hardware will enable image analysis on the fly for real-time shear flow deformation cytometry.

## Supporting information

SI Video 1: Tank Treading

## Acknowledgements

This study was supported by the Deutsche Forschungsgemeinschaft (TRR-SFB 225 subproject A01), and the European Union’s Horizon 2020 research and innovation programmes No 812772 (project Phys2BioMed, Marie Skł odowska-Curie grant) and No 953121 (project FLAMIN-GO). We thank Jonas Hazur and Aldo Boccaccini for helpful discussions and for providing the alginate.

## Methods

The measurement setup is depicted in Fig. 1a. Cells are suspended in a high-viscosity medium (e.g. a 2% alginate solution), and are pressed via a 10 cm long, 1 mm inner diameter silicone tube through a 5.8 cm long microfluidic channel with a square cross section of 200 × 200 µm (CS-10000090; Darwin Microfluidics, Paris, France). The driving air pressure of typically 1–3 bar is regulated with a pressure regulator (KPRG-114/10, Knocks Fluid-Technik, Selm, Germany) and can be switched on or off with a 3-way valve (VHK2-04F-04F; SMC, Egelsbach, Germany). The air pressure is measured with a digital pressure gauge (Digi-04 0.4%, Empeo, Germany). Cells flowing through the channel are imaged in bright-field mode at 50–500 Hz (depending on the flow speed) with a CMOS camera (acA720-520um, Basler, Germany) using a 40x 0.4 NA objective (Leica) in combination with a 0.5x video coupler attached to an inverted microscope. After passing the microchannel, the cells are collected in a waste reservoir.

### Cell culture

Cells are cultured at 37 °C, 5% CO_2_ and 95% humidity and are split every 2–3 days for up to 20 passages.

### Preparing cells for rheological measurements

Our method for measuring viscoelastic cell properties requires that the cells, if they are adherent to a cell culture dish (NIH-3T3, vim(-/-), vim(-/-)des), are brought into suspension. For cells grown in 75 cm^2^ flasks, we remove the medium and wash the cells 3 times with 10 ml of 37°C PBS. After removing the PBS, 5 ml of 0.05% trypsin/EDTA in PBS are added and distributed over the cells, and after 10 s, 4 ml of the supernatant are removed. Cells are then incubated for 3–5 min at 37°C, 5% CO_2_. 5 ml of 37°C cell culture medium (Table 1) are added to the flask, and the cells are counted. If cells are already in suspension (THP-1 and HL60 cells), the above steps are omitted. 10^6^ cells are taken out of the flask, centrifuged for 5 min at 25 rcf (NIH-3T3, vim(-/-) and vim(-/-)des) or 290 rcf (HL-60 and THP-1) to remove the supernatant, gently mixed in 1 ml of equilibrated suspension fluid (see below), transferred to a 2 ml screw-cup test tube, and centrifuged at 150 rcf for 30 seconds to remove air bubbles.

**Table 1.** cell line-specific composition of culture medium all % values are (v/v)%

| cells | Base medium | serum | PenStrep | GlutaMAX | Geneticin | Sodium pyruvate | MEM NEAA | HEPES |
| --- | --- | --- | --- | --- | --- | --- | --- | --- |
| NIH-3T3 | DMEM | 10% BCS | 1% | - | - | - | - | - |
| vim(-/-) | DMEM | 10% FCS | 1% | 1% | - | - | - | - |
| vim(-/-)des | DMEM | 10% FCS | 1% | 1% | 1 mg/ml | - | - | - |
| HL60 | RPMI | 10% FCS | 1% | - | - | - | - | - |
| THP-1 | RPMI | 10% FCS | 1% | - | - | 1 mM | 1% | 10 mM |

all % values are (v/v)%
NIH-3T3: mouse embryonic fibroblast cells (No. CRL-1658; American Type Culture Collection)
vim(-/-): Mouse embryonic fibroblast cells derived from vimentin(-/-) mice (kindly provided by Prof. Dr. T. M. Magin, University of Bonn) and further subcloned to eliminate desmin- or keratin-expressing cells, as described in<sup>27</sup>
vim(-/-)des: vim(-/-) MEFs re-expressing wild-type desmin, generated as described in<sup>28</sup>
HL60: human leukemic lymphoblast cells (No. CCL-240; American Type Culture Collection)
THP-1: monocytic cells (No. TIB-202; American Type Culture Collection)
DMEM: Dulbecco's Modified Eagle Medium (Gibco ref. 11995065) RPMI: Roswell Park Memorial Institute 1640 Medium (Gibco ref. 21875034)
PenStrep: 100x penicillin-streptomycin-glutamin solution (Gibco ref. 10378016)
GlutaMAX: L-alanine-L-glutamine supplement (Gibco ref. 35050-038)

### Suspension fluid preparation

Alginate solution is prepared freshly for the next day. Sodium alginate powder (Vivapharm alginate PH176, batch nr. 4503283839, JRS Pharma GmbH, Rosenberg, Germany, or alginic acid sodium salt from brown algae, A0682, Sigma Aldrich, for THP1 cells) is dispersed at a concentration of 1.5%, 2% or 2.5% (w/v) in serum-free cell culture medium (Table 1). The alginate solution is mixed overnight with a magnetic stirrer at room temperature until all powder has been dissolved. The suspension fluid is then equilibrated by incubating for 6 hours at 37°C, 5% CO_2_. When prepared with RPMI media (but not when prepared with DMEM nor Sigma Aldrich alginate), the alginate solution is filtered with a 0.45 µm filter before use. 1 ml of alginate solution are then added to the cell pellet of 10^6^ cells in the Falcon tube and mixed using a positive displacement pipette (15314274, Gilson/Fisher Scientific) by slowly (∼ 2 s cycle time) and repeatedly (10x) sucking the liquid in and out. The alginate-cell suspension is then transferred into a 2 ml screw-cup test tube and centrifuged for 30 seconds at 150 rcf to remove air bubbles.

### Drug treatment

Drugs are mixed in the alginate for at least 15 min at 350 rpm inside an incubator (37°C, 5% CO_2_, 95% relative humidity) prior to mixing-in the cells. Cells are prepared as described above and mixed with the alginate-drug mixture using a positive displacement pipette by slowly (∼ 2 s cycle time) and repeatedly (10x) sucking the liquid in and out. The alginate-drug-cell suspension is transferred into a 2 ml screw-cup test tube and incubated for a prescribed time at 37°C, 95% rH. Prior to measurements, the alginate-drug-cell suspension is centrifuged at 150 rcf for 30 seconds to remove air bubbles.

Inhibition of actin polymerization on NIH-3T3, vimentin-knockout and desmin-knockin MEFs is performed with cytochalasin D (Cat. No. C8273; Sigma-Aldrich, St. Louis, MO). Cytochalasin D is dissolved in DMSO at a stock concentration of 20 mM. The equilibrated alginate (3 ml) is either mixed with cytochalasin D to final concentrations of 2 µM, or mixed with DMSO to final concentration of 0.01% (DMSO control), or mixed with 3 µ l of DMEM (negative control). Cells harvested from a single cell culture flask are split into 3 groups of 10^6^ cells, each group is suspended in one of the alginate solutions as described above, stored in an incubator for 15 min (alternating between either negative control of DMSO control), 30 min (drug-treated), and 45 min (alternating between either DMSO control or negative control), and measured.

Inhibition of actin polymerization on NIH-3T3 cells is performed with latrunculin B (LatB, Cat. No. L5288; Sigma-Aldrich, St. Louis, MO, dissolved in DMSO at a stock concentration of 2 mM). We add 2 µ l of LatB (stock) or 2 µ l of DMSO to 4 ml of alginate (final concentration 1000 nM LatB, 0.2% DMSO), and mix with a magnetic stirrer at 350 rpm for 15 minutes. 1850 µ l of the alginate-drug mixture is then added to 4 ml of alginate, mixed for 15 min, and the process is repeated to obtain a dilution series with LatB concentrations of 1000, 316, 100, 32, 20, 3.2, and 1 nM. The alginate-DMSO mixture is diluted in the same way. Cells are prepared and mixed into the alginate as described above and stored at room temperature for 10 min (LatB) or 20 min (DMSO control) prior to measurements.

### Image acquisition

Typically, 10,000 images per measurement are recorded with a CMOS camera (acA720-520um, Basler, Germany) at a frame rate of 50–500 Hz with an exposure time of 30 µs. To measure the flow speed, each cell has to be recorded in at least 2 consecutive images. Therefore, the frame rate *f r* is chosen depending on the maximum flow speed *v*_max_ and the width of the region of interest (ROIx): *f r > v*_max_ / (0.5 ROIx). In our setup, the ROIx is 248 µm, resulting in a maximum flow speed of 41 mm/s for a frame rate of 500 Hz. To prevent motion blur, however, we keep the maximum flow speed to about 20 mm/s.

Fluorescent images can be acquired in parallel with the bright field images. A 300 mW diode-pumped solid-state laser (wavelength 473 nm, VA-I-N-473; Viasho, Beijing, China) serves as an epifluorescent light source, and a beam splitter projects the bright field and fluorescent images onto two synchronized cameras. To separate the light paths, the bright-field illumination is long-pass filtered (>590 nm), and a band-pass filter (500–550 nm) is placed in front of the camera for the fluorescent channel.

We provide software for image acquisition (see below under Software flow chart), which includes a live-viewer and user-friendly interface for entering meta information (e.g. applied pressure, suspension medium, drug treatments) and configuration settings (e.g. frame rate, total number of images to be stored). The software is based on the pypylon library to record the images, and Python^29^ and Qt to provide the user interface.

### Cell shape analysis

We normalize the bright-field images by subtracting the mean and dividing by the standard deviation of the pixel intensities. A neural network (U-Net^30^, tensorflow^31^) trained on labeled images of different cell types and suspension media detects the cell outline and generates a binary mask, to which an ellipse is fitted (*x,y* position of the ellipse center, its semi-major (*a*) and semi-minor axis (*b*), and the angle of orientation *β* of the major axis with respect to the flow (x) direction, see Fig. 1d,e,^32^). Binary masks that do not conform to an elliptical shape based on circumference or solidity criteria (e.g. due to cell doublets or erroneous cell outlines due to poor image contrast) are discarded.

### Finding the channel mid plane and center line

Prior to recording the images, the microscope must be precisely focused to the mid plane (z = 0, see Fig. 1) of the channel. To do so, we apply a small pressure (50–100 Pa) to the suspended cells and focus the microscope in phase contrast mode to the bottom of the microchannel, which can be unambiguously identified by stationary or very slowly flowing small debris. We then move the objective up by 75 µm, which corresponds to half the microchannel’s height (100 µm) divided by the refractive index of the suspension medium. We confirmed that the reproducibility of the method is within ±1.7 µm (rms) when a 40x 0.6 NA objective is used.

The channel center line (*y* = 0, see Fig. 1) is identified from the flow speed profile as a function of the radial (*y*) position. Flow speed is computed by tracking cells over subsequent images and dividing the distance they have moved in *x*-direction by the time difference between images. A polynomial of the form

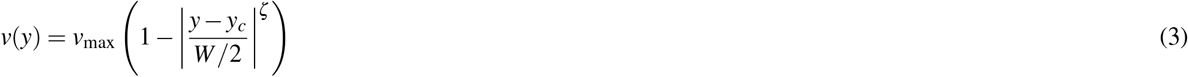

is then fitted to the velocity profile to identify the center position of the channel (*y*_*c*_), with the maximum flow speed *v*_max_ at the channel center as the second fit parameter, and the exponent *ζ* as the third fit parameter. *W* is the channel width. The fit parameter *y*_*c*_ is then used to shift the image y-coordinate origin to the channel center. This procedure ensures that the channel does not need to be precisely centered in the camera’s field of view during the measurements. However, the channel should be aligned as precisely as possible with the field of view. To ensure alignment, we recommend to rotate the camera, as opposed to the slide that holds the channels.

### Shear stress profile inside a channel with a square cross section

The fluid shear stress *σ* in the mid plane of a channel (blue shading in Fig. 1b) with length *L* and square cross section of height *H* and width *W* (Fig. 1a) only depends on the radial position *y* and the total applied pressure Δ*P* according to an infinite-series expression^3^

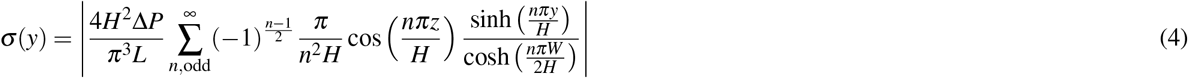

For all practical purposes, it is sufficient to compute the infinite series for the first 100 terms.

Eq. 4 assumes laminar uniaxial parallel flow and neglects entrance and exit effects, which is justified for a long and narrow channel as used in this study (*L* = 5.8 cm, *W* = *H* = 200 µm). Note that for a given channel geometry and pressure gradient Δ*P/L*, the shear stress profile *σ* (*y*) does not depend on the viscosity of the fluid. Eq. 4 remains approximately valid also for non-Newtonian e.g. shear-thinning fluids. Eq. 4 predicts that the shear stress is zero in the center of the channel and monotonically increases towards the channel wall (Fig. 1d).

Because of the finite size of a cell, the acting shear stress is calculated as the average of *σ* (*y* − *r*) and *σ* (*y* + *r*), where *y* is the center of mass of the cell, and *r* is the equivalent radius of the cell 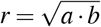, with semi-major axis *a* and semi-minor axis *b*.

### Velocity profile, shear rate profile, and viscosity

The fit function (Eq. 3) only approximates the true velocity profile, which is sufficient to efficiently and robustly find the channel center. For subsequent computations that require higher precision, we determine the velocity profile by integrating the shear rate. We compute the shear rate 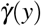 as the shear stress *σ* (Eq. 4) divided by the viscosity *η*:

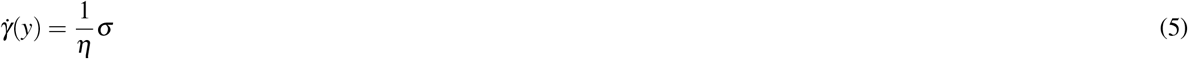

For shear thinning fluids such as alginate solutions, the viscosity *η* is not constant but depends on the shear rate 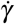. We describe the shear thinning behaviour of the viscosity by the Cross-model^33^:

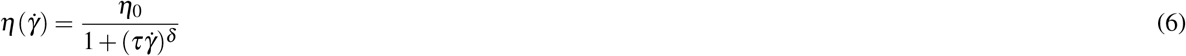

with zero-shear viscosity *η*_0_, relaxation time *τ* and power-law shear shear-thinning exponent *δ*.

When Eq. 6 is inserted into Eq. 5, we obtain

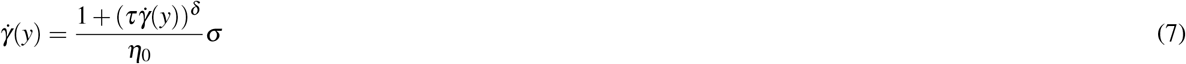

This equation can be written as

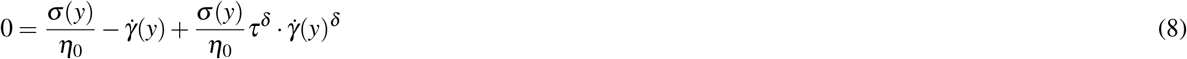

and numerically solved for 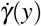 by root finding using the Newton-Raphson method.

Finally, to obtain the velocity profile *v*(*y*), we integrate the numerically obtained shear rate 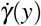 over the channel, using 5 point Gaussian quadrature:

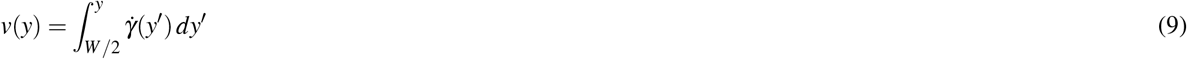

with the boundary condition *v*_*y*=*W/*2_ = 0. The viscosity parameters (*η*_0_, *τ, δ*) that best match the velocity profile are determined as follows. We choose five Gaussian quadrature points *y*^′^ between (0,*W/*2) and numerically compute 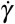 at the quadrature point *y*^′^ using Eq. 8. To ensure convergence, we start iterating with a value of 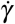 that yields the maximum of the right-hand side of Eq. 8 plus a small number *ε*. The weighted sum of 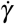 at the Gaussian quadrature points *y*^′^ is then the velocity at the radial position *y*. This procedure is repeated for different values of (*η*_0_, *τ, δ*) until a minimum of the squared differences between the measured and fitted velocity profile is found.

We find that the rheological parameters (*η*_0_, *τ, δ*) of the suspension medium obtained this way closely agree with cone-plate rheology measurements^34^. Moreover, the velocity profile for different pressure values can be accurately predicted (SI Fig. S1), demonstrating that Eq. 6 accurately describes the shear thinning behavior of the suspension fluid.

### Computing the shear strain from the cell shape

Suspended cells under zero shear stress have an approximately circular shape with radius *r*_0_. When exposed to constant shear stress, the cell deforms to an elliptical shape semi-major axis *ã* = *a*/*r*_0_ and semi-minor axes 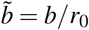 (in *x, y*-direction) and 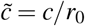 (in z-direction), normalized to the radius *r*_0_ of the undeformed cell, so that 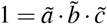. Assuming the cell consists of an incompressible material and the stress inside the deformed cell is uniform, the strain *ε* can be computed from 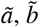 and 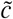 using (Eq. 10)^1^.

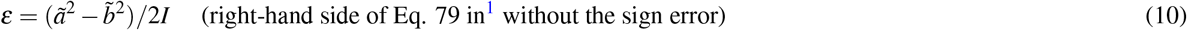

This requires solving a set of shape integrals that depend on the semi-major axis *a* and a semi-minor axis *b*.

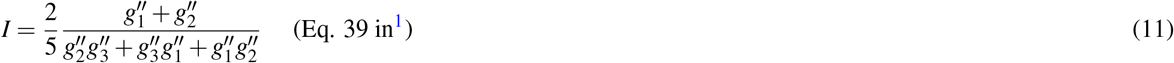

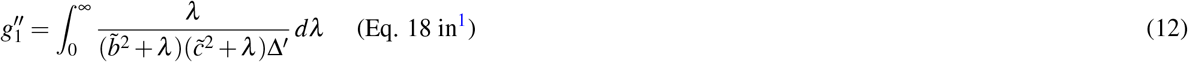

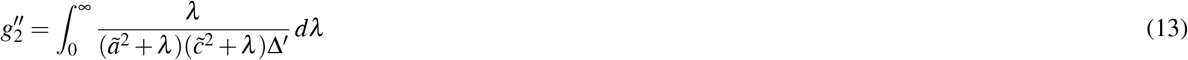

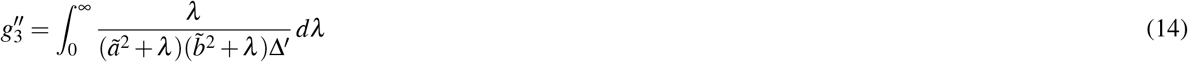

With the integration variable *λ*. Δ^′^ is defined as

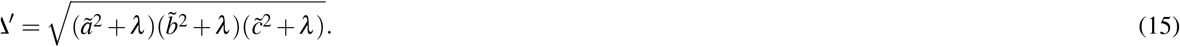

The shape integral *I* is pre-computed for different ratios of *ã* and 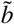 and then taken from a look-up table.

### Computing the cells’ storage and loss modulus

We calculate *G*^′^ from *σ, β, a, b* according to^1^

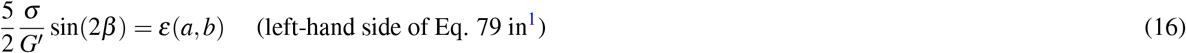

We calculate *G*^′′^ from *β, ã*, 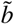, *η, ω* according to^1^

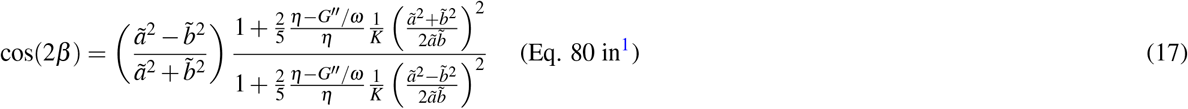

with

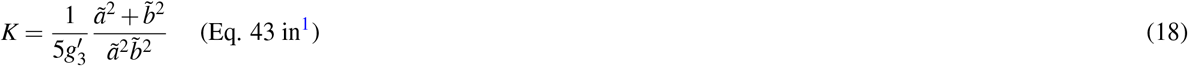

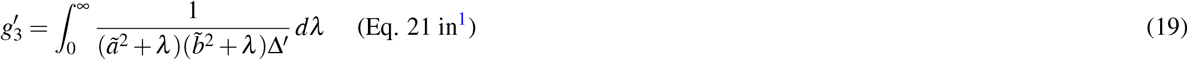

The frequency *ω* at which *G*^′^ and *G*^′′^ is obtained using Eq. 16 and 17 is the tank-treading frequency.

### Tank treading

We measure the tank-treading frequency as follows. We observe each cell as it travels through the field-of-view and cut-out small image frames with the cell at its center (Fig. 2h). We then track the movement of characteristic small features (using optical flow estimated by the TV-L1 algorithm^32,35^, and calculate their speed and distance during their rotation around the cell’s center. The speed versus the ellipse-corrected radius is fitted with a linear relationship to determine the average angular speed (Fig. 2i). The slope of this relationship is taken as the rotation frequency of the cell.

In cases where the tank-treading frequency cannot be measured (e.g. due to poor contrast or absence of cell-internal features that can be tracked), we estimate the tank-treading frequency following the approach outlined in^5^. Data shown in Fig. 2j,k demonstrate that the measured rotation rate 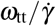 (angular frequency divided by the local shear rate) collapses onto a master relationship when plotted against the shear rate. The tank-treading frequency *ω*_tt_ of the cells can then be predicted with an empirical relationship according to

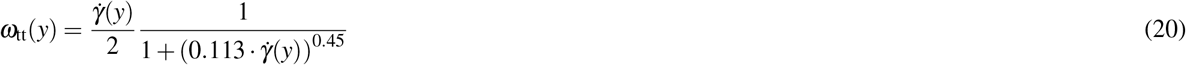

when 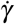 is given in units of 1/s^5^.

### Scaling the rheology

Cells show power-law rheology according to Eq. 1, which implies that the cell stiffness *k* and the power-law exponent *α* (cell fluidity) fully describe the cell rheological properties. Cell stiffness *k* and cell fluidity *α* can be obtained from *G*^′^ and *G*^′′^ measured at the tank-treading frequency *ω*_tt_ by rearranging Eq. 1 as follows:

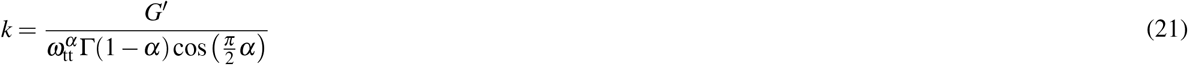

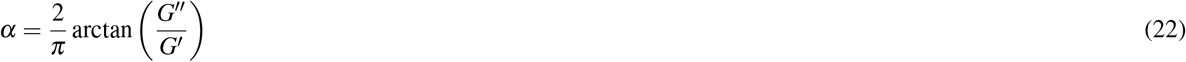

We use a Gaussian kernel density estimation^36,37^ to compute the mode of the 2-D distribution for stiffness *k* and fluidity *α*, which corresponds to the “most representative” cell with the highest joint probability for stiffness *k* and fluidity *α*.

### Software flow-chart

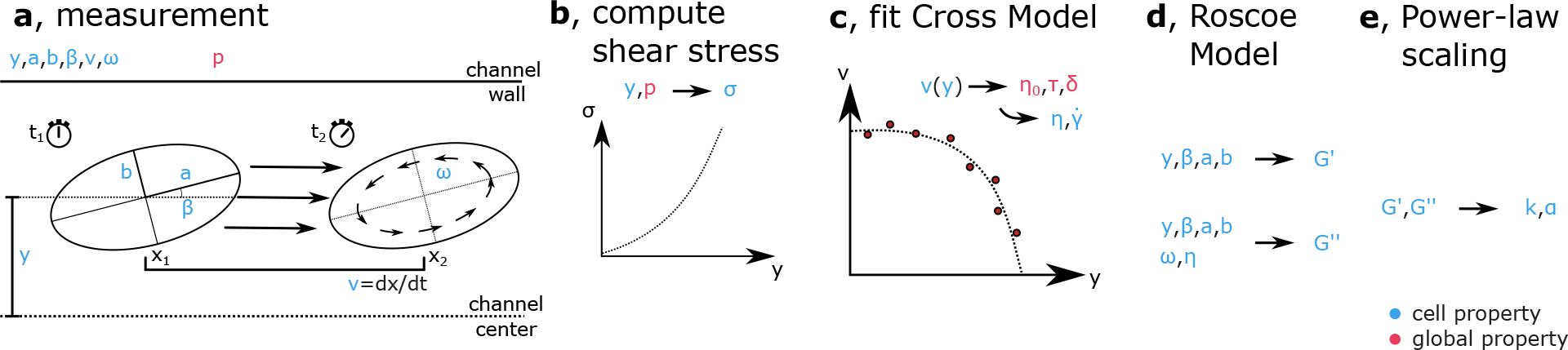

In the following, we summarize the sequence of steps and procedures for measuring cell mechanical properties with our method:

a. First, typically 10,000 image frames of cells flowing through the channel are recorded with an image acquisition program (recording.py, https://github.com/fabrylab/Deformation_Cytometer). Second, the images are analyzed off-line with an evaluation pipeline (evaluate.py, https://github.com/fabrylab/Deformation_Cytometer). The pipeline loads the images and finds and segments cells at the focal plane using a neural network^30^. From the segmented cell shape, morphological properties (*x,y* position, half major and minor axes *a* and *b*, orientation *β*, solidity, circumference) are extracted using the regionprops method of the skimage library^32^. Poorly or erroneously segmented cells that deviate from an elliptical shape are filtered out based on circumference and solidity criteria. From a measurement with 10,000 image frames, typically 5,000–10,000 cells are identified for subsequent analysis. Next, the program identifies cells that are detected across multiple subsequent frames, based on shape and position, computes the flow speed, and applies an particle image velocimetry algorithm to extract the tank treading frequency *ω*_tt_. Eq. 3 is then fitted to the speed versus y-position relationship of all cells, yielding the channel center *y*_*c*_ and the maximum flow speed *v*_max_.
b. The shear stress acting at the center position of each cell is computed using Eq. 4.
c. The shear rate at the center position of each cell is computed using a set of equations as described above (Eqs. 5–9). This procedure also yields the parameters that describe the viscosity and shear-thinning rheology of the suspension fluid (Eq. 6).
d. The cell strain is computed from the half major and minor axis *a* and *b* using Eq. 10. Subsequently, *G*^′^ and *G*^′′^ of each cell at its tank treading frequency is computed using Eq. 16 and Eq. 17.
e. To compare the mechanical properties of cells that have experienced different tank-treading frequencies, we scale *G*^′^ and *G*^′′^ to a frequency of 1 rad/s using Eq. 22 and Eq. 21, yielding the stiffness *k* and fluidity *α* of individual cells. The average stiffness *k* and fluidity *α* of the cell population is determined from the maximum of the 2-dimensional Gaussian kernel density computed using the `scipy.stats.gaussian_kde` method of the scipy library^36,37^.

### PAAm reference bead preparation

Polyacrylamide hydrogel microparticles (PAAm beads) are produced using a flow-focusing PDMS-based microfluidic chip described in^38^. Briefly, a stream of a polyacrylamide pre-gel mixture is squeezed by two counter-flowing streams of an oil solution to form droplets with a mean diameter in the range of 11.5–12.5 µm. The oil solution is prepared by dissolving ammonium Krytox surfactant (1.5% w/w), N,N,N’,N’-tetramethylethylenediamine (0.4% v/v), and acrylic acid N-hydroxysuccinimide ester (0.1% w/v) in hydrofluoroether HFE 7500 (Ionic Liquid Technology, Germany). The pre-gel mixture is obtained by dissolving and mixing acrylamide (40% w/w), bis-acrylamide (2% w/w) and ammonium persulfate (0.05% w/v) (all from Merck, Germany) in 10 mM Tris-buffer (pH 7.48). Particles with three different elasticities are obtained by diluting the pre-gel mixture in Tris-buffer to final acrylamide-bisacrylamide concentrations of 3.9%, 5.9%, 6.9% respectively. Alexa Fluor 488 Hydrazide (ThermoFisher Scientific, Germany) is dissolved in D.I. water (stock solution 3 mg/ml) and added to the mixture for a final concentration of 55 µ g/ml to make the particles fluorescent. Droplet gelation is carried out at 65 °C for 12 hours. The droplets are washed and resuspended in 1x PBS.

### Atomic force microscopy (AFM) of cells and PAAm beads

AFM-based microrheology measurements for PAAm beads are performed using a Nanowizard 4 (JPK BioAFM, Bruker Nano GmbH, Berlin). The measurements are carried out using a wedged cantilever with a flat surface parallel to the measurement dish. The cantilever is prepared by applying a UV curing glue to a tipless cantilever (PNP-TR-TL, nominal spring constant *k* = 0.08 N/m used for the stiff (6.9% *C*_AAmBis_) beads, or Nanoworld or Arrow-TL1, nominal spring constant *k* = 0.03 N/m used for the medium (5.9% *C*_AAmBis_) and soft (3.9% *C*_AAmBis_ beads) as described in^39^. Prior to each experiment, the optical lever sensitivity is measured from the force-distance relationship of a polystyrene bead attached to a glass surface, and the cantilever spring constant is measured using the thermal noise method^40^. Measured spring constants are 0.09 N/m for PNP-TR-TL cantilevers, and 0.018 N/m for Arrow-TL1cantilevers.

To perform the AFM microrheology measurements, the cantilever is lowered with a speed of 10 µm/s until a force between 1–3 nN is reached, corresponding to an indentation depth *δ* _0_ between 1.5–3 µm. The cantilever is then sinusoidally oscillated with an amplitude of 30 nm for a period of 10 cycles. This procedure is repeated for different oscillation frequencies in the range between 0.1–150 Hz. To extract the complex shear modulus *G*^∗^ of the PAAm beads, the force-indentation curves are analyzed as described in^24^ using the Hertz model that describes the deformation of a soft sphere between two flat surfaces in the limit of small deformations. The complex shear modulus is then computed according to

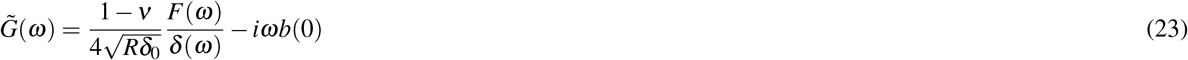

where *ν* is the Poisson ratio of the PAAm bead (assumed to be 0.5), *ω* is the angular frequency of the oscillations, *F*(*ω*) and *d*(*ω*) are the Fourier transforms of the force and indentation signal, *R* is the radius of the PAAm bead, *δ* _0_ is the initial indentation, and *b*(0) is the hydrodynamic drag coefficient of the cantilever with the surrounding liquid. The hydrodynamic drag coefficient is measured as described in^41^ and estimated to be *b*(0) = 5.28 Ns/m for PNP-TR-TL cantilevers and *b*(0) = 29.7 Ns/m for Arrow TL1 cantilevers.

AFM-based measurements for THP1 cells are performed with 4-sided regular pyramidal-tipped MLCT-bio-DC(D) cantilevers (Bruker). The spring constant of the cantilever is measured from the thermal noise spectrum in air, and the optical lever sensitivity is measured from the thermal noise spectrum in liquid^42^. The cells are immobilized to plastic petri dishes coated with poly-L-lysine at a concentration of 0.01 mg/mL for 10 minutes. Force curves are measured at 3 or more positions around the cell center for a constant indentation speed of 5 µm/s up to a maximum force of 0.8 Nn. At each position, at least 3 force-distance curves are obtained. We determine the viscoelastic step-response stress relaxation function *E*(*t*) of the cell by least-square fitting the theoretical force response to the measured force curve during indentation with a pyramidal tip^43^.

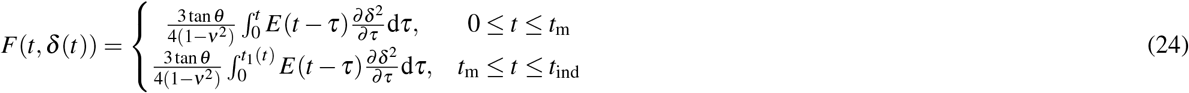

where *F* is the force acting on the cantilever tip; *δ* is the indentation depth; *t* is the time since initial contact, *t*_*m*_ is the duration of approach phase, *t*_ind_ is the duration of complete indentation cycle), and *t*_1_ is the auxiliary function determined by the equation

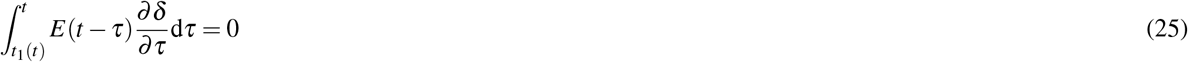

The viscoelastic step response function *E*(*t*) is assumed to follow the relationship

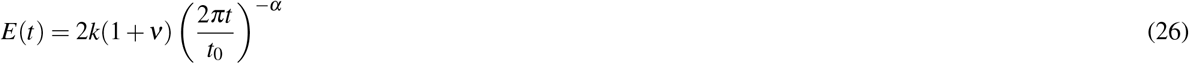

where the reference time *t*_0_ is set to 1 s so that *k* is the cell’s shear modulus measured at time *t* = 0.159 s (corresponding to *ω* = 1 rad/s as in the flow deformability measurements). The cell’s Poisson ratio *ν* is assumed to be 0.5, and *α* is the cell’s fluidity.

### Rheology of alginate solutions

We measure the viscosity of the alginate solution at a temperature of 25°C at shear rates between 0.01 s^−1^ and 1000 s^−1^ using a temperature-controlled rheometer (DHR-3, TA-Instruments, USA) with stainless steel cone and plate (diameter of 40 mm with a cone angle of 2° and a 65 µm truncation gap). Temperature is controlled with a Peltier-element. Equilibration time and measurement time are set to 30 seconds for every measurement point (logarithmic sweep, 5 points per decade). Every sample is rested for three minutes inside the rheometer to ensure temperature equilibration. A solvent trap with deionized water is used to prevent drying of the alginate samples.

### Cell cycle measurement with Fucci

We use NIH-3T3 cells that display the fluorescent ubiquitination-based cell cycle indicator (FastFUCCI) reporter system after lentiviral transduction. The lentivirus is generated by transfection of Lenti-X 293T cells (Takara, #632180) with pBOB-EF1-FastFUCCI-Puro (Addgene, #86849), a packaging plasmid psPAX2 (Addgene, #12260), and an envelope plasmid pCMV-VSV-G (Addgene, #8454), using Lipofectamine 2000 reagent (Invitrogen, #11668-019). 48 hours after transfection, infectious lentivirus-containing supernatant is harvested, centrifuged (500 × g, 10 minutes), and ten-fold concentrated using the Lenti-X-concentrator reagent (Takara, #631232). NIH-3T3 cells are seeded 24 hours prior to transduction at a density of 10 000 per cm^2^. 3 days after transduction, cells are cultured for at least 5 additional days in medium containing puromycin (5 µ g/ml) to select successfully transduced cells.

In our shear flow deformation cytometry setup, we measure only the green fluorescence signal, indicating cells in S, G2 and early M-phase^15^, and deduce that cells with a green fluorescence intensity below a certain threshold are in G1 phase. To set this threshold, we measure both the red fluorescence signal (indicating cells in G1 phase^15^) and the green fluorescence signal of individual cells prior to harvesting, using an epifluorescence microscope. We then compute the green-fluorescence intensity threshold, normalized to the median intensity that best separates the cells in G1 phase from the cells in S, G2 and early M-phase. Because some cells fluoresce green and red at the same time, 22.6% of cells in G1 phase and 2.4% of the cells in S, G2 and early M-phase are erroneously classified when the classification is based on the green fluorescence signal alone. After harvesting and suspending the cells in alginate, they are measured in the shear flow setup. Bright-field images are analyzed as described above to segment cells that are in focus, and the fluorescence intensities are averaged over the segmented cell area.

## Supplementary Information

### S1 Accuracy and predictive power of the velocity fit

**Figure S1.**
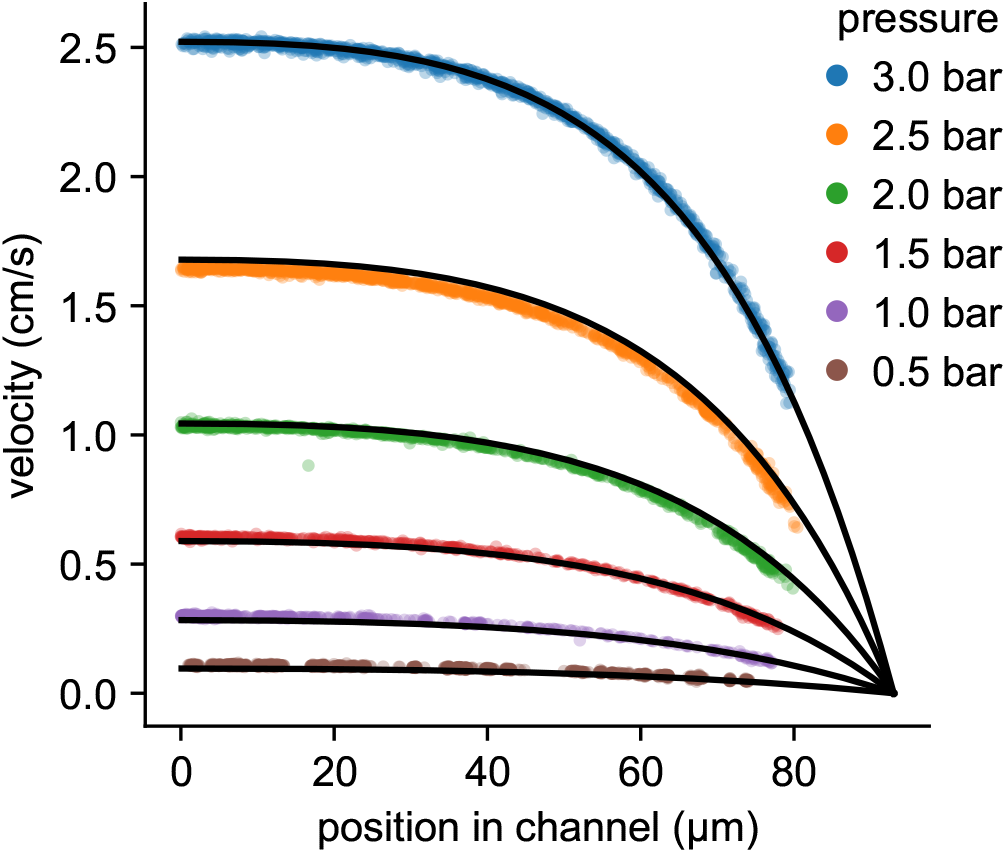
Velocity profile of a 2% alginate solution as a function of the *y*-position in the channel for different driving pressures (same data as in Fig. 2d). The top black line through the 3 bar data points shows the velocity profile fitted to the 3 bar data (each point representing the measured velocity of an NIH-3T3 cell) using Eq. 4– 9. Based on this fit, we determine the Cross-model rheological parameters (*η*_0_, *τ, β*) of the alginate solution and then predict the velocity profile for all other driving pressures (0.5–2.5 bar). The excellent agreement between measured and predicted velocities confirms the applicability of Eq. 4– 9.

### S2 Size dependence

**Figure S2.**
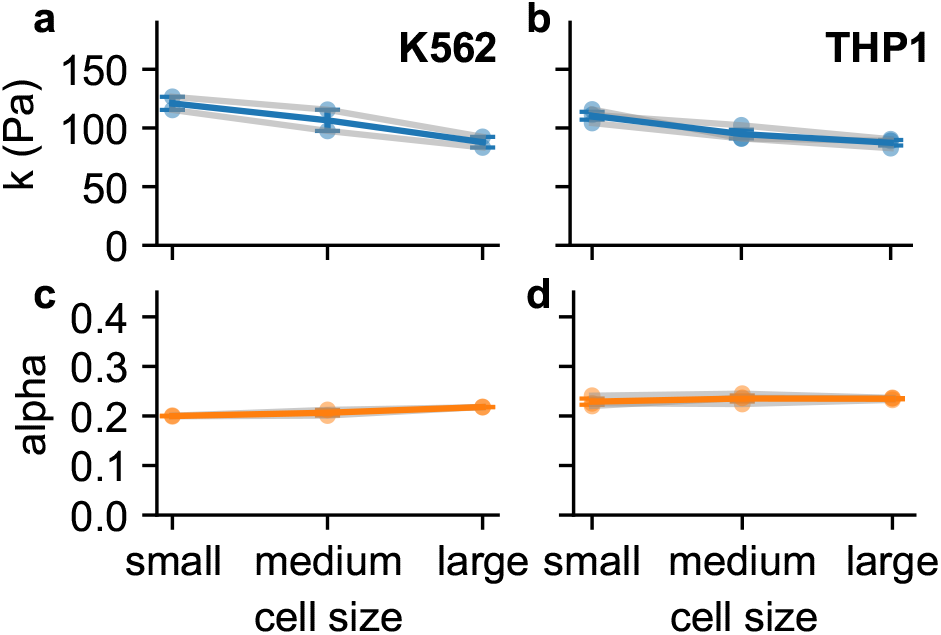
Stiffness *k* and fluidity *α* of K562 cells (2 independent measurements, measured at a pressure of 3 bar) and THP1 cells (3 measurements, 2 bar) for different cell sizes. Data a shown as mean ± se from 2 (or 3) independent measurements with n > 1500 cells for each measurement. Cells are grouped according to their size (equivalent diameter of the undeformed cell) with an equal number of cells in each group (K564 small < 6.7 µm, large > 8.4 µm; THP1 small < 9 µm, large > 10 µm).

### S3 Calibration with polyacrylamide beads

**Figure S3.**
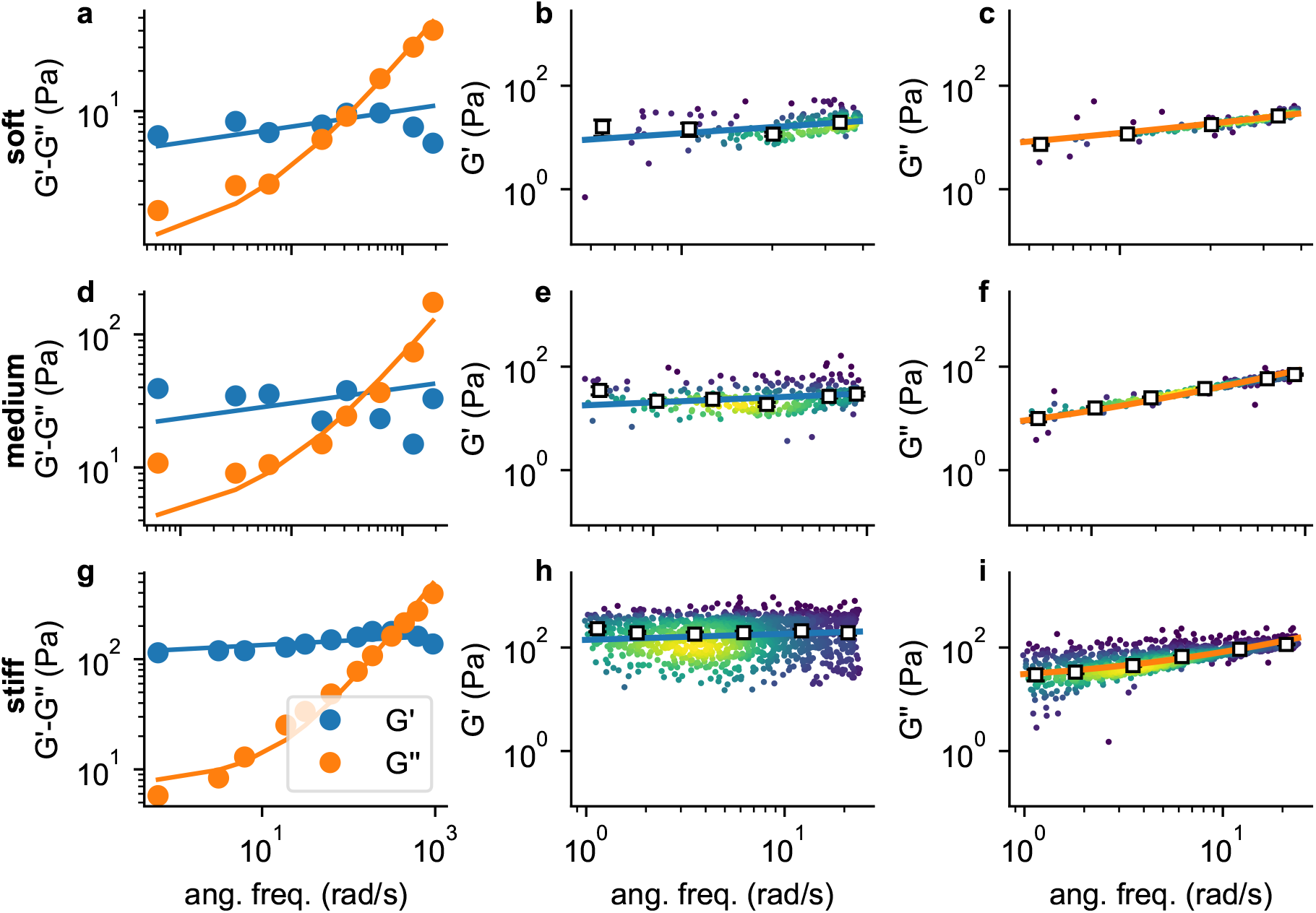
Frequency-dependent shear modulus *G*^′^ (blue) and loss modulus *G*^′′^ (orange) of polyacrylamide (PAAm) beads (soft: 3.9% *C*_AAmBis_ (top row); medium: 5.9% *C*_AAmBis_ (middle row); stiff: 6.9% *C*_AAmBis_ (bottom row)) measured with atomic force microscopy (left column) and shear flow deformation cytometry (middle and right columns). Lines show the fit of Eq. 2 to the AFM data (left column, solid circles show mean values from *n* ≥9 beads), or to the shear flow cytometry data (middle and right column, each dot represents the data from one bead, colors represent Gaussian kernel density, white squares show the median values over equal-sized bins).

## References

1. Roscoe, R. On the rheology of a suspension of viscoelastic spheres in a viscous liquid. J. Fluid Mech. 21 (1967).

2. Urbanska, M. et al. A comparison of microfluidic methods for high-throughput cell deformability measurements. Nat Methods 17, 587–593, 10.1038/s41592-020-0818-8 (2020).

3. Delplace, F. Laminar flow of newtonian liquids in ducts of rectangular cross-section an interesting model for both physics and mathematics. Int J Theor Math Phys 4, DOI:10.5923/j.ijtmp.20180802.04 (2018).

4. Fischer, T. M., Stohr-Lissen, M. & Schmid-Schonbein, H. The red cell as a fluid droplet: tank tread-like motion of the human erythrocyte membrane in shear flow. Science 202, 894–6, 10.1126/science.715448 (1978).

5. Snijkers, F. et al. Effect of viscoelasticity on the rotation of a sphere in shear flow. J. Non-Newtonian Fluid Mech. 166, 363–372, 10.1016/j.jnnfm.2011.01.004 (2011).

6. Fabry, B. et al. Scaling the microrheology of living cells. Phys Rev Lett 87, 148102. (2001).

7. Desprat, N., Richert, A., Simeon, J. & Asnacios, A. Creep function of a single living cell. Biophys J 88, 2224–33 (2005).

8. Balland, M. et al. Power laws in microrheology experiments on living cells: Comparative analysis and modeling. Phys Rev E Stat Nonlin Soft Matter Phys 74, 021911 (2006).

9. Cai, P. et al. Quantifying cell-to-cell variation in power-law rheology. Biophys J 105, 1093–102, 10.1016/j.bpj.2013.07.035 (2013).

10. Hecht, F. et al. Imaging viscoelastic properties of living cells by afm: Power-law rheology on the nanoscale. Soft Matter 11, 4553–4732 (2015).

11. Bonakdar, N. et al. Mechanical plasticity of cells. Nat Mater 15, 1090–4, 10.1038/nmat4689 (2016).

12. Smith, B. A., Tolloczko, B., Martin, J. G. & Grutter, P. Probing the viscoelastic behavior of cultured airway smooth muscle cells with atomic force microscopy: stiffening induced by contractile agonist. Biophys J 88, 2994–3007 (2005).

13. Lange, J. R. et al. Microconstriction arrays for high-throughput quantitative measurements of cell mechanical properties. Biophys J 109, 26–34, 10.1016/j.bpj.2015.05.029 (2015).

14. Kalcioglu, Z. I., Mahmoodian, R., Hu, Y. H., Suo, Z. G. & Van Vliet, K. J. From macro-to microscale poroelastic characterization of polymeric hydrogels via indentation. Soft Matter 8, 3393–3398, 10.1039/c2sm06825g (2012).

15. Sakaue-Sawano, A. et al. Visualizing spatiotemporal dynamics of multicellular cell-cycle progression. Cell 132, 487–98, 10.1016/j.cell.2007.12.033 (2008).

16. Cordes, A. et al. Prestress and area compressibility of actin cortices determine the viscoelastic response of living cells. Phys Rev Lett 125, 068101, 10.1103/PhysRevLett.125.068101 (2020).

17. Zhelev, D. V., Needham, D. & Hochmuth, R. M. Role of the membrane cortex in neutrophil deformation in small pipets. Biophys J 67, 696–705 (1994).

18. Gossett, A. J. et al. Hydrodynamic stretching of single cells for large population mechanical phenotyping. Natl. Acad. Sci. 109, 6, 10.1073/pnas.1200107109 (2012).

19. Otto, O. et al. Real-time deformability cytometry: on-the-fly cell mechanical phenotyping. Nat Methods 12, 199–202, 4 p following 202, 10.1038/nmeth.3281 (2015).

20. Fregin, B. et al. High-throughput single-cell rheology in complex samples by dynamic real-time deformability cytometry. Nat Commun 10, 415, 10.1038/s41467-019-08370-3 (2019).

21. Rowat, A. C. et al. Nuclear envelope composition determines the ability of neutrophil-type cells to passage through micron-scale constrictions. J Biol Chem 288, 8610–8, 10.1074/jbc.M112.441535 (2013).

22. Lange, J. R. et al. Unbiased high-precision cell mechanical measurements with microconstrictions. Biophys J 112, 1472–1480, 10.1016/j.bpj.2017.02.018 (2017).

23. Mietke, A. et al. Extracting cell stiffness from real-time deformability cytometry: Theory and experiment. Biophys. J. 109, 2023–2036, 10.1016/j.bpj.2015.09.006 (2015).

24. Alcaraz, J. et al. Microrheology of human lung epithelial cells measured by atomic force microscopy. Biophys J 84, 2071–9. (2003).

25. Sollich, P. Rheological constitutive equation for a model of soft glassy materials. Phys Rev E 58, 738–759 (1998).

26. Patteson, A. E., Carroll, R. J., Iwamoto, D. V. & Janmey, P. A. The vimentin cytoskeleton: when polymer physics meets cell biology. Phys. Biol. 18, Artn01100110.1088/1478-3975/Abbcc2 (2021).

27. Gregor, M. et al. Mechanosensing through focal adhesion-anchored intermediate filaments. FASEB J 28, 715–29, 10.1096/fj.13-231829 (2014).

28. Herrmann, H. et al. Dual functional states of r406w-desmin assembly complexes cause cardiomyopathy with severe intercalated disc derangement in humans and in knock-in mice. Circulation 142, 2155–2171, 10.1161/CIRCULATIONAHA.120.050218 (2020).

29. Van Rossum, G. & Drake Jr, F. L. Python reference manual (Centrum voor Wiskunde en Informatica Amsterdam, 1995).

30. Ronneberger, O., Fischer, P. & Brox, T. U-net: Convolutional networks for biomedical image segmentation. In International Conference on Medical image computing and computer-assisted intervention, 234–241 (Springer, 2015).

31. Abadi, M. et al. Tensorflow: A system for large-scale machine learning. In 12th {USENIX} Symposium on Operating Systems Design and Implementation ({OSDI} 16), 265–283 (2016).

32. van der Walt, S. et al. scikit-image: image processing in Python. PeerJ 2, e453, 10.7717/peerj.453 (2014).

33. Cross, M. M. Rheology of non-newtonian fluids: a new flow equation for pseudoplastic systems. J. colloid science 20, 417–437 (1965).

34. Muller, S. J. et al. Flow and hydrodynamic shear stress inside a printing needle during biofabrication. PLoS One 15, e0236371, 10.1371/journal.pone.0236371 (2020).

35. Zach, C., Pock, T. & Bischof, H. A duality based approach for realtime tv-l1 optical flow. In Hamprecht, F. A., Schnörr, C. & Jähne, B. (eds.) Pattern Recognition, 214–223 (Springer Berlin Heidelberg, Berlin, Heidelberg, 2007).

36. Silverman, B. W. Density estimation for statistics and data analysis (Routledge, 2018).

37. Jones, E., Oliphant, T., Peterson, P. et al. SciPy: Open source scientific tools for Python (2001–).

38. Girardo, S. et al. Standardized microgel beads as elastic cell mechanical probes. J Mater Chem B 6, 6245–6261, 10.1039/c8tb01421c (2018).

39. Stewart, M. P. et al. Wedged afm-cantilevers for parallel plate cell mechanics. Methods 60, 186–94, 10.1016/j.ymeth.2013.02.015 (2013).

40. Hutter, J. L. & Bechhoefer, J. Calibration of atomic-force microscope tips. Rev. Sci. Instruments 64(7), 1868–1873 (1993).

41. Alcaraz, J. et al. Correction of microrheological measurements of soft samples with atomic force microscopy for the hydrodynamic drag on the cantilever. Langmuir 18(3), 716–721 (2002).

42. Sumbul, F., Hassanpour, N., Rodriguez-Ramos, J. & Rico, F. One-step calibration of afm in liquid. Front. Phys. 8, 301 (2020).

43. Efremov, Y. M., Wang, W. H., Hardy, S. D., Geahlen, R. L. & Raman, A. Measuring nanoscale viscoelastic parameters of cells directly from afm force-displacement curves. Sci Rep 7, 1541, 10.1038/s41598-017-01784-3 (2017).

